# A Grain of Salt

**DOI:** 10.1101/2020.01.23.916742

**Authors:** Kelly Houston, Jiaen Qiu, Stefanie Wege, Maria Hrmova, Helena Oakey, Yue Qu, Pauline Smith, Apriadi Situmorang, Malcolm Macaulay, Paulina Flis, Micha Bayer, Stuart Roy, Claire Halpin, Joanne Russell, Caitlin Byrt, Matt Gilliham, David E. Salt, Robbie Waugh

## Abstract

We quantified grain sodium (Na^+^) content across a barley GWAS panel grown under optimal conditions. We identified a strong association with a region containing two low and one high Na^+^ accumulating haplotypes of a Class 1 HIGH-AFFINITY POTASSIUM TRANSPORTER (HKT1;5) known to be involved in regulating plant Na^+^ homeostasis. The haplotypes exhibited an average 1.8-fold difference in grain Na^+^ content. We show that an L189P substitution disrupts Na^+^ transport in the high Na^+^ lines, disturbs the plasma membrane localisation typical of HKT1;5 and induces a conformational change in the protein predicted to compromise function. Under NaCl stress, lines containing P189 accumulate high levels of Na^+^, but show no significant difference in biomass. P189 increases in frequency from wild-species to elite cultivars leading us to speculate that the compromised haplotype is undergoing directional selection possibly due to the value of Na^+^ as a functional nutrient in non-saline environments.

## Introduction

Sodium (Na^+^) is a non-essential but functional nutrient for plant growth and development^1^. This means that while plants can grow and reproduce in the absence of Na^+^, when Na^+^ is present it generally provides a range of benefits. While halophytes thrive on high Na^+^ containing soils^2^, for glycophytes, including our major cereal crops, Na^+^ becomes toxic when present above certain species-specific threshold levels. Intriguingly, many crops, including barley, have been shown to benefit from intermediate (non-toxic) levels of Na^+^, a situation that is particularly evident when levels of K^+^ in the soil are low^1, 3–10^. In such cases, Na^+^ appears capable of substituting for many of the essential roles that K^+^ ions play in plant nutrition, including enzyme activation and osmoregulation. Indeed, extensive historical evidence supports a requirement for non-toxic levels of Na^+^ to achieve maximal biomass growth in a wide range of plants^11^. It is thus somewhat ironic that, despite the demonstrated positive attributes of Na^+^, by far the majority of studies in the more recent literature focus on the negative impacts of Na^+^ (i.e. salinity) on plant growth^12, 13^. These latter investigations generally seek to explore the possible mechanisms that explain how tolerance to excess Na^+^ can be achieved, and commonly revolve around Na^+^ exclusion from the transpiration stream via active removal in the root, the partitioning of excess Na^+^ into the vacuole in Na^+^ sensitive photosynthetic tissues of the shoot, or the energy balance associated with active tolerance mechanisms^14, 15^. While understanding how to enable crops to grow more efficiently in the expanding saline environments across the globe is highly relevant, it remains important to note the majority of temperate cereal crop production is actually achieved on non-saline soils. Given the demonstrated benefits of Na^+^ as a functional nutrient, here we have taken a combined genetic and functional approach to explore the extent and causes of natural variation in Na^+^ content in barley grown in non-saline soils. Our data lead us to speculate that high Na^+^ accumulation may be a positive trait in the non-saline conditions typical of high production agricultural environments.

## Results

### Barley grain Na+ content is genetically controlled

We used Inductively Coupled Plasma Mass Spectrometry (ICP-MS) to quantify sodium (Na^+^) and potassium (K^+^) content of whole grain samples from five biological and five technical replicates of a small barley GWAS panel comprised of 131 elite 2-row spring genotypes. All plants were grown under optimal, non-saline conditions. We observed an approximate six-fold variation in grain Na^+^ (16.07 to 98.76 ppm) and greater than two-fold variation in K^+^ (2459 to 5562 ppm) contents (**Figure 1a and Supplementary Table 1**). We then used barley 50k iSelect SNP genotypic data collected from all 131 genotypes to conduct GWAS^16^. For grain Na^+^ content we observed a single highly significant association on the bottom of chromosome 4HL with a -log10(p) = 11.8, and an R^2^ = 0.45 (**Figure 1b**) consistent with prior genetic analyses of shoot Na^+^ content^2–4, 5 17–20^. The moderately high R^2^ value implies other loci and mechanisms are also involved in this trait. No significant genetic association was observed for grain K^+^ content at this locus (**Figure 1c**). The significantly associated region spanned approximately 6.6Mb, from 638,211,825 nt to 644,818,273 nt on the barley 4H physical map^21^ and contained 247 gene models (**Supplementary Table 2**). Common SNPs between the 50k and 9K iSelect platforms^22^ aligned this association with a region recently shown to contain *HvHKT1;5*^20, 23^, and in the barley genome sequence HORVU4Hr1G087960, 140kb from the top scoring SNP (chr4H_638774955), was annotated as a homolog of *OsHKT1;5.* Considering this and previous functional studies^24–27^we conclude that HORVU4Hr1G087960 is the Na^+^ specific transporter *HvHKT1;5*.

**Figure 1.**
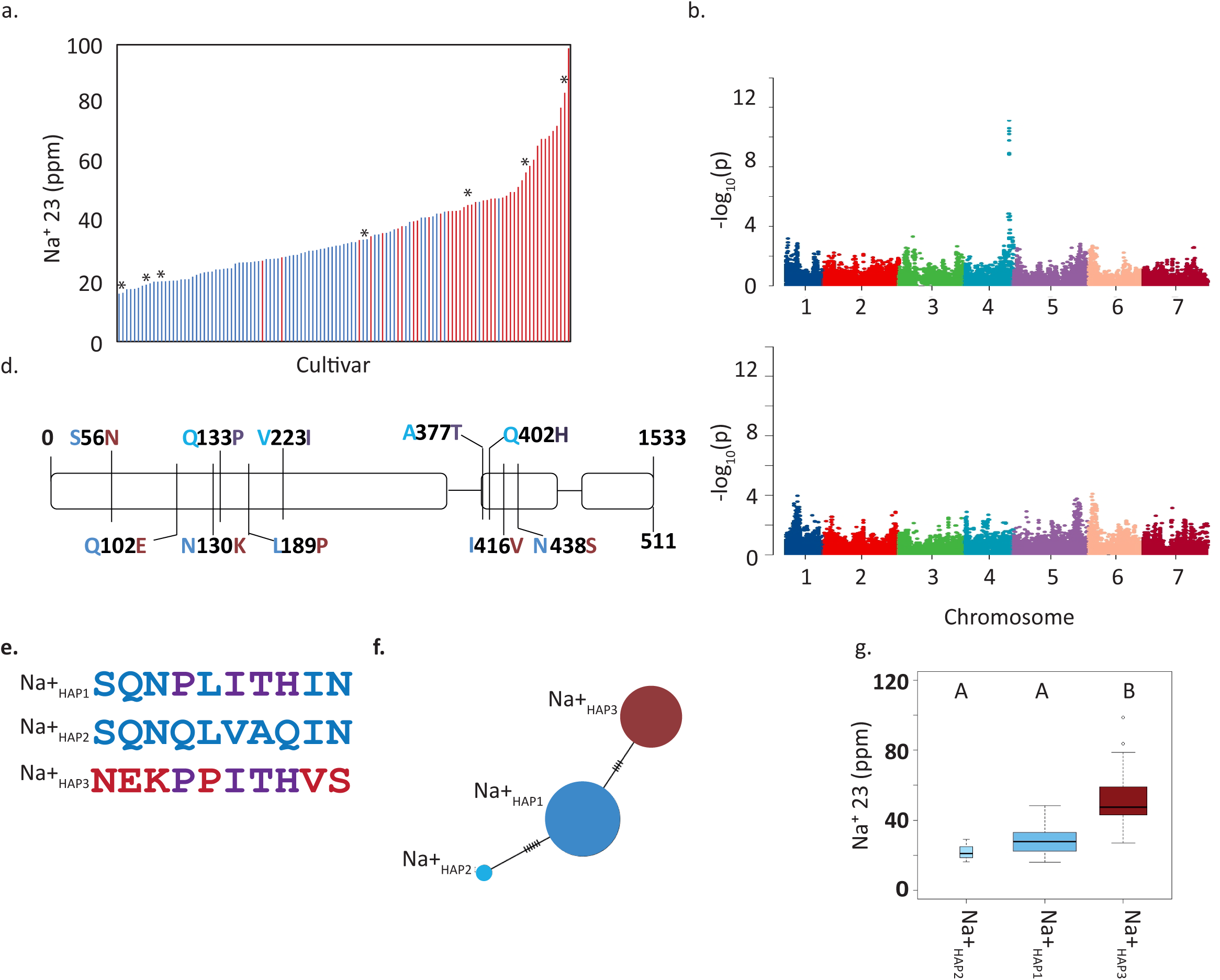
*HvHKT1;5* haplotypes influence grain Na^+^ content. **a.** Frequency distribution of BLUPs of grain sodium content quantified using ICP-MS. Blue bars represent lines containing L189 and red bars represent genotypes containing P189. Asterix indicates lines from the panel included in salt stress experiment. **b.** Manhattan plot of GWAS of grain Na^+^ 23 content, FDR threshold = -log 10(P) 6.02. **c**. Manhattan plot of GWAS of grain K^+^ content using, FDR threshold = -log 10(P) 6.02. **d**. Gene structure and nonsynonymous polymorphisms coloured according to haplotype shown in e. **e.** Haplotype summary based on non-synonymous SNPs. **f.** Haplotype analysis of non-synonymous SNPs. Circles scaled to number of individuals sharing the haplotype, and short lines represent number of SNPs differentiating haplotype groups. **g.** Box plots of grain Na^+^ contents in the three haplotypes. Centre line represents the median, boxes represent the upper and lower quartiles, whiskers extending from the box represent the 1.5x interquartile range, and circles represent outliers. The boxes are drawn with widths proportional to the square-roots of the number of observations in the groups. Different letters above boxes indicate significant difference in grain sodium content p<0.05, same letters indicate no significant difference using this threshold.

As *HvHKT1;5* is a clear candidate for causing the observed phenotypic variation we PCR-sequenced this gene from all 131 genotypes included in the GWAS. We observed 10 nonsynonymous SNPs that defined three haplotypes (defined as Na^+^_HAP1,_ Na^+^_HAP2_ and Na^+^_HAP3_) (**Figure 1d, e, f**). Na^+^_HAP1_ and Na^+^_HAP2_ correspond exactly to the HGB haplotype very recently described^5^. In our GWAS panel there was no association between *HvHKT1;5* haplotype and population structure. We observed a significant difference in mean grain Na^+^ content between genotypes containing the low grain Na^+^ haplotypes, Na^+^_HAP1_ (M=28.6, SD±72) and Na^+^_HAP2_ (M=22.2, SD ± 6.48), and the high grain Na^+^_HAP3_ allele (M=51.6, SD± 15.2); t (52)=9.07, p= 2.66E-12 (two tailed test)) (**Figure 1g**). There was no significant difference between Na^+^_HAP1_ and Na^+^_HAP2_ (M=28.6, SD± 7.72, M=22.2, SD± 6.48); t (2) =1.66, p=0.23(two tailed test)). Lines containing Na^+^_HAP3_ contain an average increase in grain Na^+^ content of 1.8-fold over Na^+^_HAP1_ and Na^+^_HAP2_. Of the 10 non-synonymous SNPs, six in complete linkage disequilibrium (LD) altered amino acid residues (S56N, Q102E, N130K, L189P, I416V, N438S) that differentiated the two low Na^+^ haplotypes (Na^+^_HAP1_ and Na^+^_HAP2_) from the high Na^+^ haplotype (Na^+^_HAP3_). The remaining four (P133Q, I223V, T377A and H402Q) differentiated low Na^+^_HAP1_ and high Na^+^_HAP3_ from Na^+^_HAP2_ (**Figure 1d, e**).

### Transcript abundance varies between *HvHKT1;5* haplotypes

As variation in transcript abundance among *HvHKT1;5* haplotypes has previously been implicated in determining variation in shoot Na^+^ content^20^, we selected genotypes that were representative of Na^+^_HAP1_ (*cv.* Golden Promise), Na^+^_HAP2_ (*cv.* Viivi) and Na^+^_HAP3_ (*cv.* Morex) and quantified *HvHKT1;5* transcript abundance by qRT-PCR in a range of tissues after growth without added Na^+^(**Figure 2**). We observed significant differences (P<0.05) in the normalised gene expression in each haplotype across tissues. In general terms, *HvHKT1;5* was more highly expressed in roots compared to shoots, and low Na^+^_HAP1_ and Na^+^ _HAP2_ were expressed more highly than high Na^+^_HAP3_. The highest overall expression was observed in the maturation zone of the roots in *cv.* Viivi (Na^+^_HAP2_) with *in situ* hybridisations using sections from this region showing that *HvHKT1;5* was predominantly expressed in the xylem parenchyma and endodermal cells adjacent to the xylem vessels (**Figure 2c**)^24, 25, 28^. These results are consistent with *HvHKT1;5* transcript abundance in root and shoot tissues reflecting the observed high and low grain Na^+^ haplotypes.

**Figure 2.**
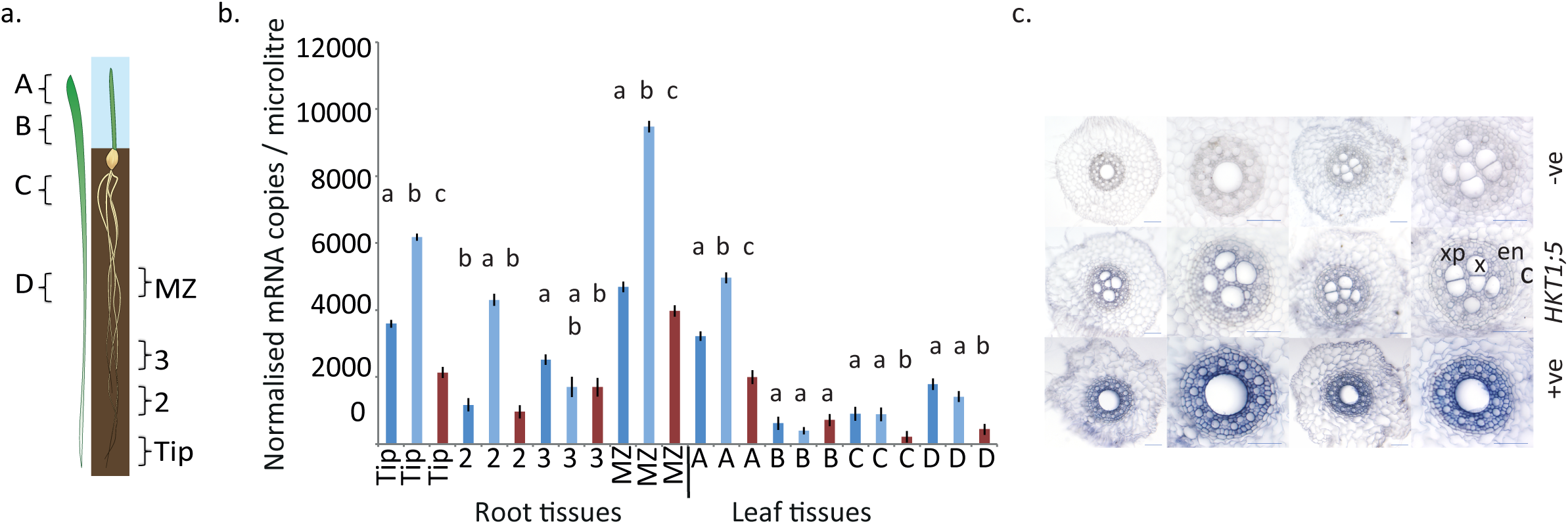
Expression of *HvHKT1;5* haplotypes. **a.** Sections used for quantification of transcript abundance (after Burton *et al*., 2004^46^). **b**. Quantification of *HvHKT1;5* in root (Tip, 2, 3, and MZ) and shoot (A, B, C, and D) tissues from 12-day old barley plants. Error bars represent SE. Dark blue - Na^+^_HAP1_ (*cv.* Golden Promise), pale blue =Na^+^_HAP2_ (*cv.* Vivii), red=Na^+^_HAP3_ (*cv.* Morex) differences from after carrying out an ANOVA followed by Tukey HSD within tissue type, same letters in lower case indicate no significant difference using P <0.05. **c**. *In-situ* localisation of *HvHKT1;5* in 2-week-old barley root tissue from *cv* Golden Promise, Na^+^_HAP1_ (grown without added NaCl). Top: negative controls with no RT (reverse transcription), Middle *HvHKT1;5* with c, cortex; en, endodermis; x, xylem; xp, xylem parenchyma labelled in red. Bottom *Hv18S* rRNA (positive control). Scale bars, 100 μm.

### A single amino acid substitution in HvHLT1;5_HAP3_ disrupts Na^+^ transport function

Despite the current lack of evidence for natural functional variation in HvHKT1;5^20^ we were interested in testing whether the observed haplotypes influenced *in vivo* Na^+^ transport properties. We assembled and independently tested constructs expressing Na^+^_HAP1_, Na^+^_HAP2_ or Na^+^_HAP3_ in *Xenopus laevis* oocytes using two-electrode voltage-clamp (TEVC) experiments. Oocytes injected with cRNA of Na^+^_HAP1_ (or Na^+^_HAP2_, data not shown) showed significant inward currents in the presence of external Na^+^ but not K^+^ (**Figure 3**) consistent with *HvHKT1;5* being a Na^+^-specific transport protein. With the external Na^+^ concentration increased from 1 mM to 30 mM, a two-fold increase in Na^+^ conductance was observed (**Figure 3**). For Na^+^_HAP3_, Na^+^ and K^+^ elicited currents were similar to water-injected controls with the conductance unaltered when external Na^+^ concentration was increased (**Figure 3**), indicating that Na^+^_HAP3_ was severely compromised in its ability to transport Na^+^ across the plasma membrane.

**Figure 3.**
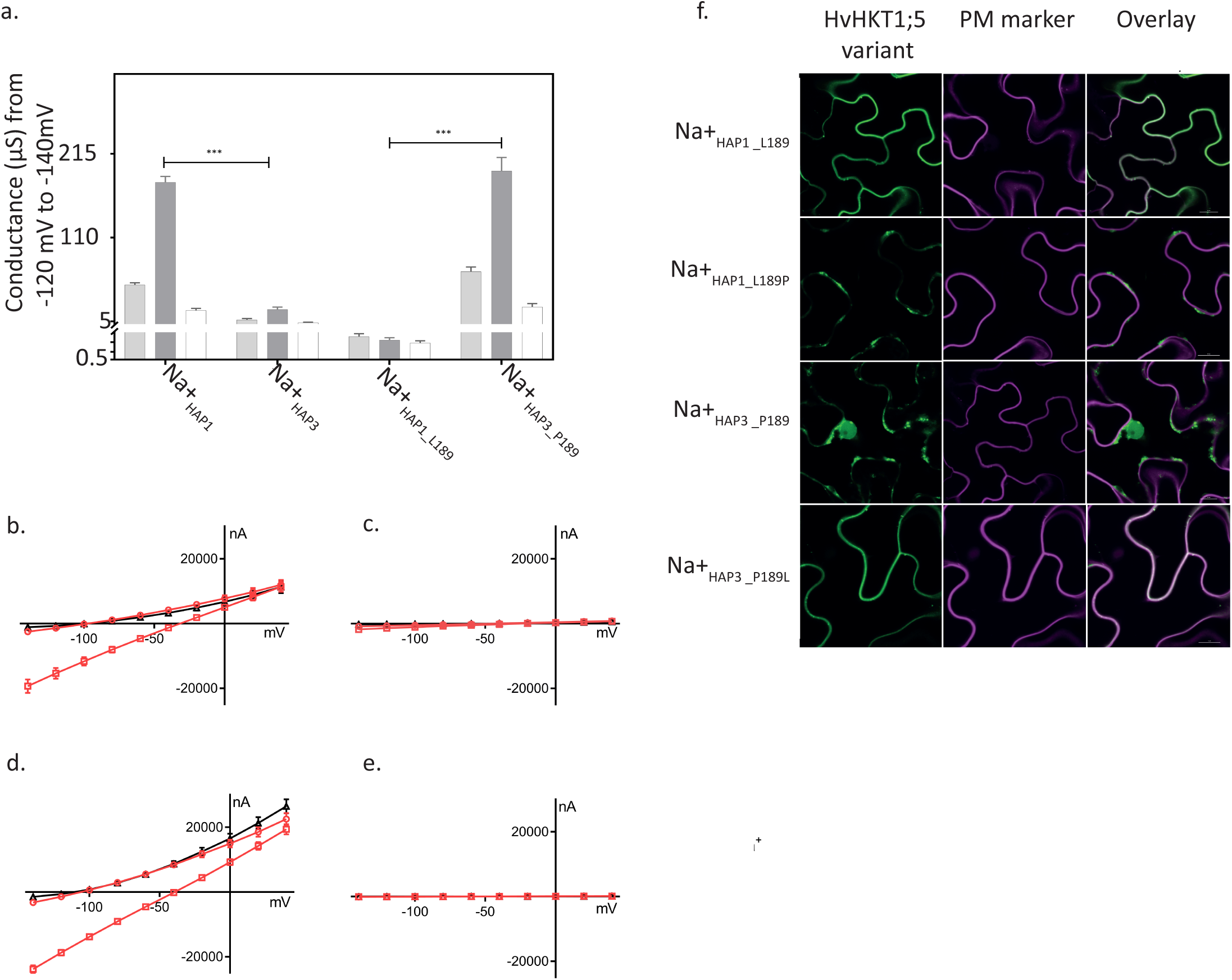
Heterologous expression of HvHKT1;5 variants in *Xenopus laevis* oocytes and *Nicotiana benthamiana* epidermal cells. **a**. Cation (Na^+^ and K^+^) conductance (−140 mV to −120mV) from HvHKT1;5 haplotypes cRNA-injected oocytes perfused with 1mM (light grey boxes), 30 mM Na^+^ (dark grey boxes) and 30 mM K^+^ (white boxes). Data are means ± SEM of currents, n = 11–21 (***P < 0.001), combined from 3 independent experiments. **b.- e.** Representative I-V curves of cRNA from HvHKT1;5 haplotypes injected into *Xenopus* oocytes (n=4) clamped at −140 mV to 40 mV in Na+ or K+ solutions. Red circles (1mM Na+), red squares (30 mM Na+), black triangles (30 mM K+). **b.** Na^+^_HAP1___L189_; **c.** Na^+^_HAP3___P189_; **d.** Na^+^_HAP3___L189_; **e.** Na^+^_HAP3___P189_. **f.** Transient co-expression of GFP-HvHKT1;5 variants with CBL1n-RFP plasma membrane marker in *Nicotiana benthamiana* leaf epidermal cells. GFP signal in the left panel (green), RFP-signal in the middle (magenta), overlay on the right (co-localisation of green and magenta signals appears in white). Scale bars = 10 μm

Publicly available data for HKT1;5 led us to focus on four single amino acid residue changes as potentially causal for compromised transport (N57S, P189L, V416I and S438N) ^20, 28^. We swapped these candidate amino acids residues individually into the compromised high Na^+^_HAP3_ and quantified their impact on Na^+^ conductance by TEVC in the oocyte system. In comparison to high Na^+^_HAP3,_ Na^+^_HAP3_L189_ showed levels of Na^+^-dependent conductance that were comparable to low Na^+^_HAP1_ (**Figure 3**). The reciprocal substitution (P189) into low Na^+^_HAP1_ showed significantly reduced Na^+^ conductance, comparable to high Na^+^_HAP3_ (**Figure 3**). No other substitution converted a low Na^+^ haplotype to a high Na^+^ haplotype or *vice versa*; however, the Na^+^_HAP1_V416_, reduced but did not abolish the Na^+^-dependent conductance of Na^+^_HAP1_ (**Supplementary Figure 1**). These data support the conclusion that the naturally occurring P189 amino acid residue in Na^+^_HAP3_ compromises the function of HvHKT1;5, and that certain variants (e.g. Na^+^_HAP1_V416_) can also affect Na^+^ dependent conductance in TEVC experiments.

### HvHLT1;5_HAP3_ does not localise to the plasma membrane

Consistent with their role in Na^+^ retrieval from the xylem sap, HKT1;5 proteins have been previously shown to localise specifically to the plasma membrane (PM). We were therefore interested in whether the observed structural and functional variation had consequences for HvHKT1;5 subcellular localisation. We transiently co-expressed N-terminally GFP-tagged HvHKT1;5_HAP3___L189P_ variants with a plasma membrane (PM)-marker in *Nicotiana benthamiana* epidermal cells. Confocal imaging revealed that the low Na^+^ variant HvHKT1;5_HAP3_L189_ was almost exclusively localised at the PM (**Figure 3**). However, the high Na^+^ HvHKT1;5_HAP3_P189_ did not co-localise with the PM-marker; the GFP-signal was instead localised to internal cell structures (**Figure 3**). Introduction of P189 into HvHKT1;5_HAP1_ phenocopied the GFP-signal pattern of cells transformed with HvHKT1;5_HAP3_ (**Figure 3**). This GFP-signal pattern in cells expressing HvHKT1;5 haplotypes harbouring P189 may suggest protein degradation.

To explore this further we constructed 3D molecular models of HvHKT1;5_HAP3_L189_ and HvHKT1;5_HAP3_P189_, in complex with Na^+^ using the *B. subtilis* KtrB K^+^ transporter (Protein Data Bank genotype 4J7C, chain I) as a template with K^+^ substituted by Na^+^^29, 30^ (**Supplementary Figure 2**). In the structural models, detailed analysis of the micro-environments around α-helices 4 and 5 revealed that the α-helix 4 of low Na^+^ allele HvHKT1;5_HAP3_L189_ established a network of four polar contacts at separations between 2.7 Å to 3.1 Å with A185, V186, Y192 and S193 neighbouring residues. However, these were not formed in high Na^+^ HvHKT1;5_HAP3_P189_, which only established two polar contacts at separations between 2.5 Å to 2.7 Å with S193. We observed a positive correlation between the structural characteristics of α-helices 4 and 5 (trends in angles based on α-helical planes), differences in Gibbs free energies of forward (P189L) and reverse (L189P) mutations, and the ability to produce Na^+^ fluxes across oocyte membranes. Combined with our previous observations we hypothesise that P189 in HvHKT1;5 does affect protein structure, potentially triggering protein degradation and/or insertion into the plasma membrane thereby reducing Na^+^ retrieval from the xylem with bulk flow ultimately elevating Na^+^ in the grain.

### The impact of *HvHKT1;5* haplotypes under salt stress

While our original data were collected from plants grown under optimal conditions, most recent reports in the literature focus on the impact of variation at HKT1;5 on natural tolerance to growth in saline environments^31–33^. We therefore explored the impact of *HvHKT1;5* variants on a range of phenotypic traits after growth in 0mM, 150mM and 250mM added NaCl **(Figure 4a)**. While we observed confounding between allele, haplotype and line (see full analysis and statistics given in **Supplementary dataset 1)** we can nevertheless conclude that grain Na^+^ content is influenced significantly by both allele (L189P) and treatment (NaCl); lines containing the functionally compromised P189 allele accumulate higher levels of Na^+^ than lines containing L189 and show a larger difference between control and salt treatments. All lines with L189 accumulate less Na^+^ in the grain than those with P189, with one genotype, Maris Mink, having especially high grain Na^+^. This line had the second highest levels of grain Na^+^ when grown as part of the GWAS panel (**Supplementary Table 1**). Despite the higher Na^+^ content, no clearly detrimental effect of P189 was observed on biomass yield (**Figure 4b**). 250mM NaCl had a strong and consistent negative influence on total biomass across all lines. There was no influence of L189P on grain K^+^ content (P>0.05), although there was an effect of treatment on this trait (P = 0.02) (**Supplementary Figure 3).** As previous studies of HKT1;5s generally focus on shoot Na^+^, we also examined shoots from the same plants. Again, we observed that an interaction between allele and treatment significantly influenced leaf Na^+^ content (P<0.001), with lines containing L189 accumulating less Na^+^ in leaf tissue than those with P189 (**Supplementary Figure 4, Supplementary Table 5**), mirroring our observations in grain.

**Figure 4.**
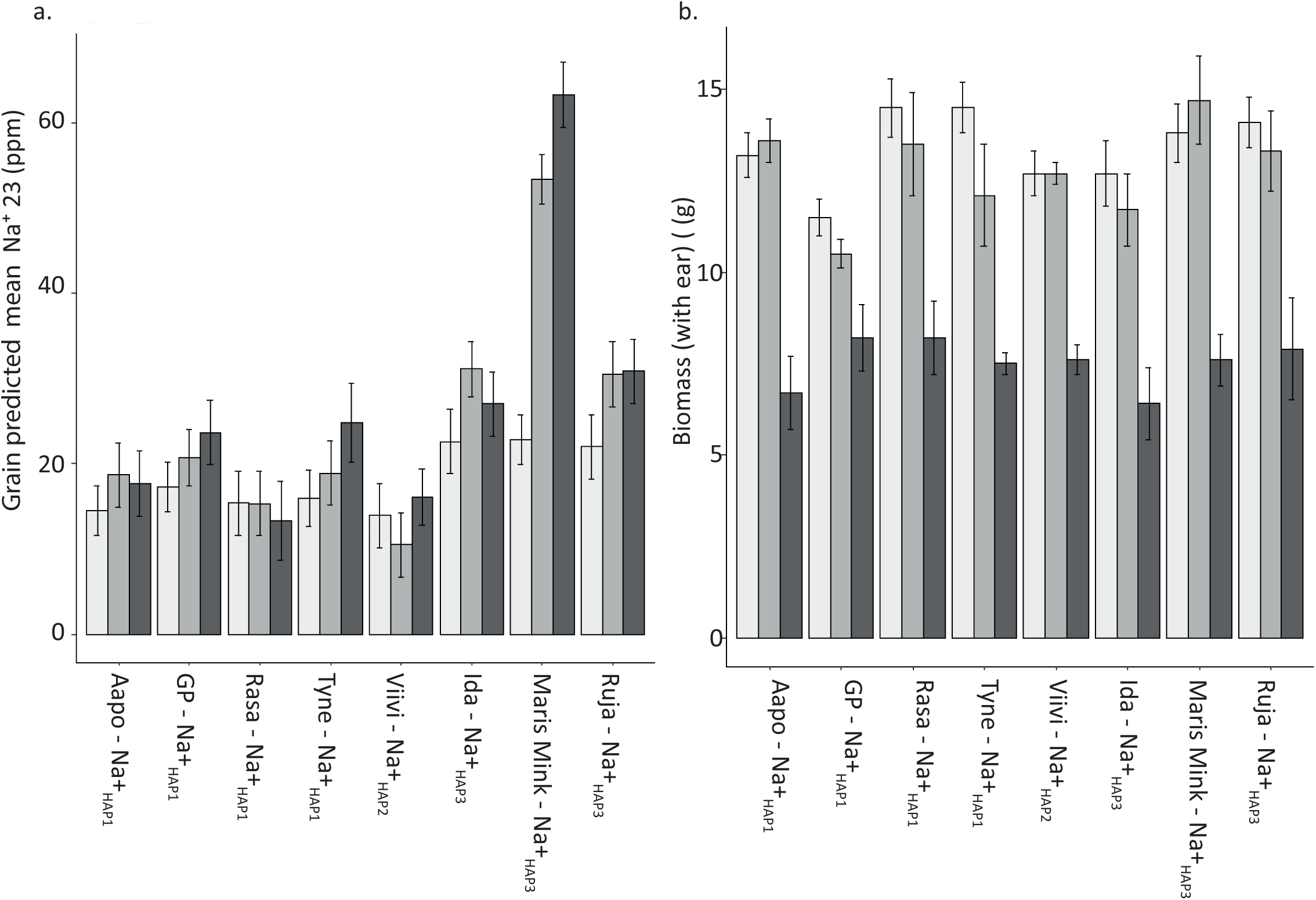
Influence of L189P polymorphism in HvHKT1;5 on grain Na^+^ accumulation and other morphological traits. **a.** Predicted mean values of mature grain Na+ content (back-transformed to original scale after analysis) of barley genotypes exposed to 0mM, 150mM and 250mM NaCl at the fourth leaf stage of development. **b.** Above ground biomass (including the ear of grain) after barley genotypes were exposed to different levels of NaCl at the fourth leaf stage of development. For both **a.** and **b.** *HvHKT1;5* haplotype is shown alongside name of genotype, low grain Na^+^_HAP1,_ Na^+^_HAP2,_ and high grain Na^+^_HAP3._

### *HvHKT1;5_HAP3_* frequency increases significantly in elite NW European barley

Previously observed associations between *AtHKT1* alleles and the environment^34^ prompted us to explore whether barley *HvHKT1;5* haplotypes had any obvious evolutionary or ecological significance. We identified and downloaded orthologs of *HKT1;5* and aligned the retrieved protein sequences with MUSCLE^35^. Based on the available sequences only barley genotypes contained the L189P substitution in HKT1;5 despite comparing amino acid sequences of 11 different species **(Supplementary Figure 5)**. To explore the origin and distribution of the P189 variant, we then PCR-sequenced *HvHKT1;5* from a collection of 73 georeferenced wild barley (*H. spontaneum*) genotypes from the fertile crescent^36^. This revealed 19 additional non-synonymous SNPs defining 27 haplotypes (**Supplementary Table 3**) that are distinct from those observed in the GWAS panel, with the exception of Na^+^_HAP3_ that was found in one genotype, FT064, originating from southern Israel (latitude = 31.35, longitude = 35.12) (**Supplementary Figure 6, 7**). A neighbour joining tree based on nonsynonymous SNPs in these *H. spontaneum* genotypes plus those in the elite cultivated barley genotypes revealed three discreet clades each containing one elite line haplotype (**Supplementary Figure 7**). We then genotyped the L189P polymorphism in 184 georeferenced landraces and found that 7 genotypes (<4%), mostly located in western Europe, contained the P189 substitution (**Supplementary Figure 6, 7**). Strikingly, this frequency increased to 35% in the 131 elite genotypes used for GWAS (**Supplementary Figure 7**). This increase, which occurs across all branches of the cultivated genepool is symptomatic of what would be expected for a locus that is currently undergoing directional selection.

## Discussion

Sodium content is a complex trait that can have serious implications for plant performance and survival. By combining high density SNP-array and ionomic data collected from the grain of plants grown under non-saline conditions, we identified haplotypes of *HvHKT1;5* as the major genetic factor determining Na^+^ content in contemporary 2-row spring barley. *HvHKT1;5* has previously been implicated in conferring a degree of salinity tolerance in wheat and barley through a Na^+^ exclusion mechanism. In high Na^+^ accumulating genotypes we found that a single SNP generating an L189P amino acid residue substitution led to severely compromised HvHKT1;5 function. We propose this is likely due to a combination of protein structural changes leading to misfolding, aberrant subcellular localisation and subsequent degradation, and is compounded by higher transcript abundance of the functional alleles^20^. Although a recent study^37^identified SNPs that cause TaHKT1;5 to become non-functional the protein still localised to the plasma membrane unlike HvHKT1;5_HAP3_.

After growing representatives of each haplotype under saline conditions, our observations align with previous findings in barley^19, 20^, and in rice^25^ in response to short-term NaCl stress. However, in the latter, longer term stress (21 days of 40mM or 80 mM NaCl) led to a 72% decrease in biomass in an *OsHKT1;5* expression mutant compared to wild type. Consistent with these findings, when Munns *et al.*^31^ backcrossed a functional *Nax2* locus (*TmHKT1;5A*) from *T. monococcum* into commercial Durum wheat they observed a yield increase of 25% compared to the control when grown on saline soils. Both studies parallel the relationship observed between Arabidopsis *AtHKT1* allele and seed number from plants grown under saline conditions; wild-type Arabidopsis Col-0 produced seeds when exposed to moderate salt stress while an *athkt1* knockout mutant was virtually sterile^32^. Together they suggest that HKT1’s are critically important for maintaining fitness under saline conditions. Our data, supported by recent evidence that shoot Na^+^ accumulation in certain bread wheat genotypes is not negatively associated with plant salinity tolerance^38, 39^, question this conclusion for barley. Here we show that biomass yield (and by inference photosynthesis) is maintained in lines such as Maris Mink (HvHKT1;5_HAP3_) that have levels of grain (and leaf) Na^+^ content that would be expected to significantly impact yield. We conclude that alternative or additional mechanisms must be involved in Na^+^ tolerance in barley and that Na^+^ exclusion by HvHKT1;5 may be a relatively minor player. This discrepancy could potentially arise because, unlike barley, Arabidopsis, rice and Durum wheat are all particularly sensitive to saline conditions that may point to fundamental differences in the roles of HKT1;5 between salt tolerant and salt sensitive plants. Recently, *HvHKT1;5* RNAi knockdown lines generated in *cv.*GP (Na^+^_HAP1_L189_), were shown to exhibit an increase in shoot biomass compared to WT at increasing NaCl levels^40^. Somewhat controversially, the authors hypothesise that *HvHKT1:5* translocates Na^+^ from the root to the shoot, and the increase in biomass in the RNAi lines is due to a reduction in translocation of Na^+^ due to decreased expression of *HvHKT1;5*. However, the data presented here, in another recent study of *HvHKT1;5*^5^, and reports in several different crop species ^10, 11, 12^ indicate that the presence of a functionally compromised alleles of *HvHKT1;5* leads to an increase in shoot and grain sodium content due to a reduced ability to exclude Na^+^ from the plant.

Our observed increase in the frequency of high Na^+^ *HvHKT1;5_HAP3_* in breeding germplasm from NW Europe returns us to the possible role of Na^+^ as a functional micronutrient in agriculture and its beneficial effects on plant growth and development, particularly in low K^+^ environments. While the NW European growing environment is largely devoid of saline soils it is subject to periodic spring/summer droughts and depleting levels of soil K^+^ due to annual offtake from high yielding varieties being greater than the amount of K^+^ applied. In such situations the higher Na^+^ content of *HvHKT1;5_HAP3_* could theoretically provide a physiological advantage, for example through use of Na^+^ as a substitute for K^+^ in a range of metabolic functions or as a free and abundant osmolyte to reduce leaf water potential, increase or maintain photosynthesis and ultimately impact yield. If correct, we would be tempted to speculate that the increase in frequency may reflect ongoing positive selection during breeding for *HvHKT1;5_HAP3_* due to it providing a selective advantage. However, given it is not yet near fixation in elite genotypes, distinguishing this hypothesis from alternatives such as selection due to linkage disequilibrium with another positive trait clearly remains to be tested.

Overall we can conclude that natural allelic variation at *HvHKT1;5* has a strong influence over grain and shoot Na^+^ homeostasis in barley in non-saline and saline environments. A single SNP causing an L189P amino acid substitution likely results in a change in protein structure, that leads to aberrant sub-cellular localisation, loss of capacity to transport Na^+^ (in frog oocytes) and consequent reduction in capacity to remove Na^+^ from the transpiration stream leading to elevated levels in the shoots and grain. Of note, when grown under salt stress conditions we observed no negative consequences of the high levels of Na^+^ in the compromised haplotype on a range of life history traits, most notably biomass. Consequently, this fails to provide support for *HvHKT1;5* driven Na^+^ exclusion in the root as playing the dominant role in salinity tolerance in barley. Curiously, the identical amino acid residue substitution (L190P) was recently identified in the bread wheat genotype, Mocho de Espiga Branca, that similarly exhibits atypically high shoot Na^+^ content and significantly reduced Na^+^ conductance in TEVC experiments^39^, providing a striking example of parallel evolution in two economically important species with potential value in plant breeding.

## Methods

### Phenotypic charecterisation of grain sodium in contemporary UK genotypes for GWAS

A collection of 131 contemporary European 2-rowed spring barley genotypes, for the purposes of this study our GWAS panel, were grown in a polytunnel in Dundee, Scotland, using standard barley soil and growth conditions.

We screened the grain sodium concentration of the GWAS panel using Inductively Coupled Plasma Mass Spectrometry (ICP-MS). Barley grains were transferred into Pyrex test tubes (single grain per tube) and weighted. Samples were pre-digested overnight at room temperature with 1 mL trace metal grade nitric acid Primar Plus (Fisher Chemicals) spiked with indium internal standard followed by digestion in dry block heaters (DigiPREP MS, SCP Science; QMX Laboratories, Essex, UK) at 115°C for 4 hours. Then, 1 mL of hydrogen peroxide (Primar, for trace metal analysis, Fisher Chemicals) was added and samples were digested in dry block heater at 115°C for 2h. After cooling down, the digests were diluted to 10 mL with 18.2 MΩcm Milli-Q Direct water (Merck Millipore) and elemental analysis was performed using PerkinElmer NexION 2000 ICP-MS equipped with Elemental Scientific Inc. autosampler, in the collision mode (He). Twenty-one elements (Li, B, Na, Mg, P, S, K, Ca, Cr, Mn, Fe, Co, Ni, Cu, Zn, As, Rb, Sr, Mo, Cd and Pb) were monitored. The isotopes 23 and 39 were measured for Na and K, respectively. Liquid reference material composed of the pooled digested samples was prepared before the beginning of the sample run and was used throughout the whole samples run. It was run after every ninth sample in all ICP-MS sample sets to correct for variation between and within ICP-MS analysis runs. The calibration standards (with indium internal standard and blanks) were prepared from single element standards solutions (Inorganic Ventures; Essex Scientific Laboratory Supplies Ltd, Essex, UK). Sample concentrations were calculated using external calibration methods within the instrument software. Further data processing was performed in Microsoft Excel. For each genotype Na was measured in five biological reps, and this was repeated per five technical replicates. From these data BLUPs were predicted using GenStat (15^th^ edition).

### DNA extraction, 50k iSelect genotyping, and GWAS

DNA from 7-day old leaves was extracted for all genotypes using the QIAamp kit (Qiagen) on the QIAcube HT (Qiagen) using default settings. All samples were genotyped using the 50k iSelect SNP array as described in^16^. GWAS was carried out on adjusted variety means using the EMMA algorithm and a kinship matrix derived using Van raden in GAPIT^41^ with R version 3.5.2^42^. We anchored regions of the genome which were significantly associated with Na^+^ content to the physical map of the barley sequence to provide annotations for genes within these regions^21^. Linkage disequilibrium (LD) was calculated for regions of the genome containing significant associations between pairs of markers using a sliding window of 500 markers and a threshold of R2<0.2 using Tassel v5^43^ to allow us to identify local blocks of LD, facilitating a more precise delimitation of QTL regions. We anchored regions of the genome containing markers that passed the FDR to the physical map and then expanded this region using local LD derived from genome wide LD analysis as described above.

### Resequencing *HvHKT1;5* and sequence alignment

For the 131 genotypes of the GWAS panel we PCR amplified and Sanger sequenced the coding sequence of *HvHKT1;5*. We used the primers listed in **Supplementary Table 4** to resequence this gene. DNA was amplified and cleaned up prior to Sanger sequencing on an ABI3100 capillary sequencer using reaction mixes and conditions described^44^. Sequences were aligned in Geneious version 9.0.2(Biomatters Ltd). Haplotype networks were produced using PopART version 1.7^45^. Orthologs of HvHKT1;5 were identified using the blastx function at NCBI, and the sequences retrieved aligned in Geneious version 9.0.2 (Biomatters Ltd) using MUSCLE with default settings.

### RNA extraction and cDNA synthesis

Materials detailed in **Figure 2a** were sampled, snap frozen and stored at −80 °C for RNA extraction. The root tissue was ground to fine powder on 2010 Geno/Grinder® (SPEX SamplePrep) at 1200 RPM for 30 seconds, and RNA was extracted from the tissue powder by using Direct-Zol RNA MiniPrep (Zymo Research) according to the manufacturer’s protocol. Final elution was performed with 40 µL DNA/RNAase-Free water supplied with the kit and the eluted RNA was subsequently quantified using ND-1000 Spectrophotometer (NanoDrop Technologies). cDNA synthesis was then performed on 500ng RNA by using High Capacity cDNA Reverse Transcription Kit (Thermo Fisher Scientific) according to the manufacturer’s instruction in a 20 µL reaction and stored at −20 °C until use.

### RNA extraction and qPCR of *HvHKT1;5*

Root and shoot tissue from 12-day old roots and shoots were collected in sections as described^46^ from genotypes grown in the same polytunnel as described above, in the same conditions for RNA extraction. Each of the 4 Biological reps consisted of tissue collected from 15 individual plants. cDNA was synthesised using RNA to cDNA EcoDry™ Premix (Double Primed) (Takara) using standard conditions and used for qPCR. qPCR and the analysis of the subsequent data was carried out as described^46^ using 3 housekeeping genes, α – tubulin, GAPDH, and HSP70. Primer sequences and annealing temperatures are provided in **Supplementary Table 4.**

### Characterisation of diversity of *HKT1;5*

Species orthologs of HKT1;5 were identified using the blastx function at NCBI, and the protein sequences retrieved were aligned in Geneious version 9.0.2 (Biomatters Ltd) using MUSCLE with default settings. For the 73 *H. spontaneum* genotypes, DNA was extracted, *HvHKT1;5* amplified and Sanger sequenced as described above. For the landraces, the L189P SNP was typed in 184 georeferenced genotypes^36^ using primer pair 3 in **Supplementary Table 4** and PCR-sequencing conditions described above. The geolocation data for these genotypes is available^36^.

### *In-situ* PCR

Barley roots *in situ* PCR was followed by Athman *et al.*^47^ with the following modifications. Root cross sections (from maturation zone) were 60 μm obtained using Vibrating Microtome 7000 Model 7000smz-2 (Campden Instruments Ltd.). Thermocycling conditions for the PCR were: initial denaturation at 98 °C for 30 seconds, 35 cycles of 98 °C for 10 seconds, 59 °C (for *HvHKT1;5*) or 57 °C (for *Hv18S*) for 30 seconds, 72 °C for 10 seconds, and a final extension at 72 °C for 10 minutes. Gene specific primers for *HvHKT1;5* and *Hv18S* (positive control) are shown in **Supplementary Table 4**.

### Characterisation of HKT1;5 in oocytes

Methods for functional characterisation of HvHKT1;5 variants in *Xenopus laevis* oocytes were as described previously^30, 31, 48^. Haplotype and engineered variants of HvHKT1;5 were synthesised by GenScript (Piscataway, NJ, USA) and fragments were inserted into a gateway enabled pGEMHE vector. Nucleotides encoding HvHKT1;5 N57S, P189L, V416I and S438N were modified by site-directed mutagenesis PCR using Phusion^®^High-Fidelity DNA Polymerase (New England Biolabs, Massachusetts, USA). pGEMHE constructs were linearized using sbfI (New England Biolabs, Massachusetts, USA) followed by ethanol precipitation. Complimentary RNA (cRNA) was transcribed using using the Ambion mMESSAGE mMACHINE kit (Life Technologies, Carlsbad, CA, USA), 23 ng of cRNA (in 46 nL) or equal volumes of RNA-free water were injected into oocytes, followed by an incubation in ND96 for 24-48 h before recording. Membrane currents were recorded in the HMg solution (6 mM MgCl_2_, 1.8 mM CaCl_2_, 10 mM MES and pH 6.5 adjusted with a TRIS base) ± Na^+^ glutamate and/or K^+^ glutamate as indicated. All solution osmolarities were adjusted using mannitol at 220-240 mOsmol kg^-1^ ^31, 48^

### Transient expression of *HvHKT1*;5 in *Nicotiana benthamiana*

Transient expression of fluorescent fusion proteins was performed as described in detail^26^. In brief, *HKT1;5* coding sequences were recombined into pMDC43 to generate N-terminally GFP-tagged proteins. For co-localisation studies, nCBL1-RFP was used as a PM-marker^49^. All constructs were transformed into *Agrobacterium tumefaciens* strain Agl-1. Agroinfiltration was performed on fully expanded leaves of 4- to 6-week-old *Nicotiana benthamiana* plants. After two days, leaf sections were imaged using a Nikon A1R Confocal Laser-Scanning Microscope equipped with a 633-water objective lens and NIS-Elements C software (Nikon Corporation). Excitation/emission conditions were GFP (488 nm/500–550 nm) and RFP (561 nm/570–620 nm).

### Construction of 3D molecular models of HvHKT1;5_HAP3_L189_ and HvHKT1;5_HAP3_P189_ in complex with Na^+^

The most suitable template for cereal HKT1;5 transporter proteins was the *B. subtilis* KtrB K^+^ transporter (Protein Data Bank genotype 4J7C, chain I)^14^ as previously identified^15^. In KtrB, K^+^ was substituted by Na^+^ during modelling of all HKT1;5 proteins. 3D models of HvHKT1;5_HAP3_L189_ and HvHKT1;5_HAP3_P189_ in complex with Na^+^ were generated in Modeller 9v19^50^ as described previously^21, 52^ incorporating Na^+^ ionic radii^15^ taken from the CHARMM force field^53^, on the Linux station running the Ubuntu 12.04 operating system. Best scoring models (from an ensemble of 50) were selected based on the combination of Modeller Objective Function^54^ and Discrete Optimised Protein Energy term^55^ PROCHECK^56^, ProSa 2003^57^ and FoldX^58^. Structural images were generated in the PyMOL Molecular Graphics System V1.8.2.0 (Schrődinger LLC, Portland, OR, USA). Calculations of angles between selected α-helices in HvHKT1;5 models were executed in Chimera^59^ and evaluations of differences (ΔΔG = ΔGmut-ΔGwt) in Gibbs free energies was performed with FoldX^58^. Sequence conservation patterns were analysed with ConSurf^60, 61^ based on 3D models of HvHKT1;5 transporters.

Evaluations of stereo-chemical parameters indicated that the template and HvHKT1;5 models had satisfactory parameters as indicated by Ramachandran plots with two residues positioned in disallowed regions, corresponding to 0.5% of all residues, except of G and P. Average G-factors (measures of correctness of dihedral angles and main-chain covalent bonds) of the template, and HvHKT1;5_HAP3_L189_ and HvHKT1;5_HAP3_P189_ models, calculated by PROCHECK (0.06, −0.07 and −0.07, respectively), and ProSa 2003 z-scores (measures of C^β^-C^β^ pair interactions of −9.0, −5.6 and −6, respectively), indicated that template and modelled structures had favourable conformational energies.

### Plant material and growth conditions

Eight barley genotypes (*Hordeum vulgare*) with variation in the key functional SNP (P189L) in *HvHKT1;5* were grown for screening leaf and grain Na^+^ accumulation under salt stress conditions. This set of genotypes consist of at least three representative barley genotypes of each allele of P189L in HvHKT1;5 characterised in the elite germplasm within this study, *cv*. Golden Promise (GP) (Na^+^_HAP1_L189_), Aapo (Na^+^ _HAP1_L189_), Rasa (Na^+^_HAP1_L189_) and Tyne (Na^+^_HAP1_L189_), Viivi (Na^+^_HAP2_L189_), Ida (Na^+^_HAP3_P189_), Maris Mink (Na^+^_HAP3_P189_), Ruja (Na^+^_HAP3_P189_). Three germinated seeds from each genotype were sown in a 10 x 10 cm pot filled with a standard cereal compost mix as described above. Eight replicates were sown per genotype in a randomized design. Every 24 pots were randomized in a plastic gravel tray (56 cm x 40 cm x 4 cm), and in total 12 trays were placed in the glasshouse under long-day conditions (light:dark, 16h:8h, 18°C:14°C). At sowing, the soil moisture and weight of all pots ranged between 31-35 % (w/w) and between ∼380-385g. The three seedlings in each pot were thinned to one at the emergence of 2^nd^ leaf.

Before applying the salt treatment soil moisture content in all pots was controlled to around 25% (w/w) to provide larger NaCl uptake capacity. At the emergence of the 4^th^ leaf, a single salt treatment (150 or 250 mM NaCl) was applied directly into each tray in a 2L volume. The same volume of water was added to the control trays. The fully expanded 5^th^ leaf was harvested for Na^+^ content analysis using ICP-MS as described above to evaluate the strength of salt treatment **(Supplementary Table 5)**. Plants were harvested at maturity and the Na^+^ contents of grains and 5^th^ leaves of five of the eight replicates quantified by ICP-MS. Detailed description of the analysis of the resulting data is included in **Supplementary Data 1**.

## Data Availability

All sequences of HvHKT1;5 generated in this study are available from NCBI, accession numbers are provided in **Supplementary Table 1**.

## Acknowledgements

RW and MM acknowledge support from ERC project 669182 ‘SHUFFLE’ to RW. KH, PS, MB, JR, and RW acknowledge the Rural & Environment Science & Analytical Services Division of the Scottish

Government. CB, JQ and YQ acknowledge support from Rutherford Fund Strategic Partner Grants 2018 - Award Reference: RF-2018-30 to CH and RW. SR acknowledges financial assistance from The Australian Research Council Industrial Transformation Research Hub for Wheat in a Hot and Dry Climate (IH130200027), the Grains Research and Development Corporation (ACP00009) and The Waite Research Institute, University of Adelaide. CB acknowledges support from the Grains Research and Development Corporation (GRDC) and the Australian Research Council (FT180100476). We are thankful to the Australian Research Council for funding through CE140100008 to M.G. (J.Q., Y.Q.), FT180100476 to C.S.B. and DE160100804 to S.W. (A.S.). DES acknowledges support from BBSRC Grant BB/L000113/1.

## Author Contributions

RW, DES, MG, CB, KH, JQ, JR and SR designed experiments. KH, JQ, SW, YQ, PS, AS, MM, PF carried out experiments. KH, MH, HO, MB analysed data. The manuscript was written by KH and RW with contributions from all other authors.

## Ethics Declarations

The authors declare no competing interests.

## Supplementary Information

### Figures

**Supplementary Figure 1:** Current-voltage (I/V) curve of X. laevis oocytes expressing allelic variants of HvHKT1;5.

**Supplementary Figure 2:** Molecular models of HvHKT1;5_HAP3_L189_ and HvHKT1;5_HAP3_P189_ transporters in complex with Na^+^.

**Supplementary Figure 3:** Influence of L189P polymorphism in HvHKT1;5 on grain K^+^ accumulation.

**Supplementary Figure 4:** Influence of L189P polymorphism in HvHKT1;5 on grain Na^+^ accumulation.

**Supplementary Figure 5:** Multiple alignment of species HKT orthologues.

**Supplementary Figure 6:** Geographical distribution of L189P in barley germplasm.

**Supplementary Figure 7:** Dendrograms illustrating distribution of L189P in a wide range of barley germplasm.

### Tables (see associated Excel sheets)

**Supplementary Table 1:** Elite 2-row spring genotypes included in GWAS and sequenced for *HvHKT1;5*.

**Supplementary Table 2:** Gene models in region identified on 4H as being significantly associated with grain Na^+^ content.

**Supplementary Table 3:** *H spontaneum* and *H. vulgare* landrace *HvHKT1;5* genotypic data.

**Supplementary Table 4:** Primers used for sanger sequencing, qPCR and In-situs.

**Supplementary Table 5:** Na^+^ and K^+^ contents of 5th leaf material from 0mM, 150mM and 250mM NaCl treated plants.

### Supplementary Dataset

**Supplementary dataset:** Influence of growth in 0mM, 150mM and 250mM on a range of phenotypic traits: Full description of analytical methods.

