## Supplementary data for "A Grain of Salt"

**Contents:**

***Figures***

**Supplementary Figure 1:**	Current-voltage (I/V) curve of X. laevis oocytes expressing allelic	variants of HvHKT1;5.

**Supplementary Figure 2:**	Molecular models of HvHKT1;5HAP3_L189 and HvHKT1;5HAP3_P189	transporters in complex with Na+.

**Supplementary Figure 3:**	Influence of L189P polymorphism in HvHKT1;5 on grain K+	accumulation.

**Supplementary Figure 4:**	Influence of L189P polymorphism in HvHKT1;5 on grain Na+

accumulation.

**Supplementary Figure 5:**	Multiple alignment of species HKT orthologues.

**Supplementary Figure 6:**	Geographical distribution of L189P in barley germplasm.

**Supplementary Figure 7:**	Dendrograms illustrating distribution of L189P in a wide range of	barley germplasm.

***Tables (see associated Excel sheets*)**

**Supplementary Table 1:**	Elite 2-row spring cultivars included in GWAS and sequenced

for *HvHKT1;5.*

**Supplementary Table 2:**	Gene models in region identified on 4H as being significantly	associated with grain Na^+^ content.

**Supplementary Table 3:**	*H spontaneum* and *H. vulgare* landrace *HvHKT1;5* genotypic data.

**Supplementary Table 4:**	Primers used for sanger sequencing, qPCR and In-situs.

**Supplementary Table 5:**	Na^+^ and K^+^ contents of 5th leaf material from 0mM, 150mM and

250mM NaCl treated plants.

***Supplementary Dataset***

**Supplementary dataset:**	Influence of growth in 0mM, 150mM and 250mM on a range of phenotypic traits: Analytical methods.

**Supplementary Figures.**

**Supplementary Figure 1.** **Current-voltage (I/V) curve of X. laevis oocytes expressing allelic variants of HvHKT1;5**. **A**, High Na+ allele HvHKT1;5N57S. **B**, High Na+ allele HvHKT1;5V416I. **C,** High Na+ allele HvHKT1;5S438N. **D,** Low Na+ allele HvHKT1;5I416V. Currents were recorded in 1 mM, 30 mM of Na+ or 30 mM K+ glutamate; data represented in mean ± SEM. n= 3-6.

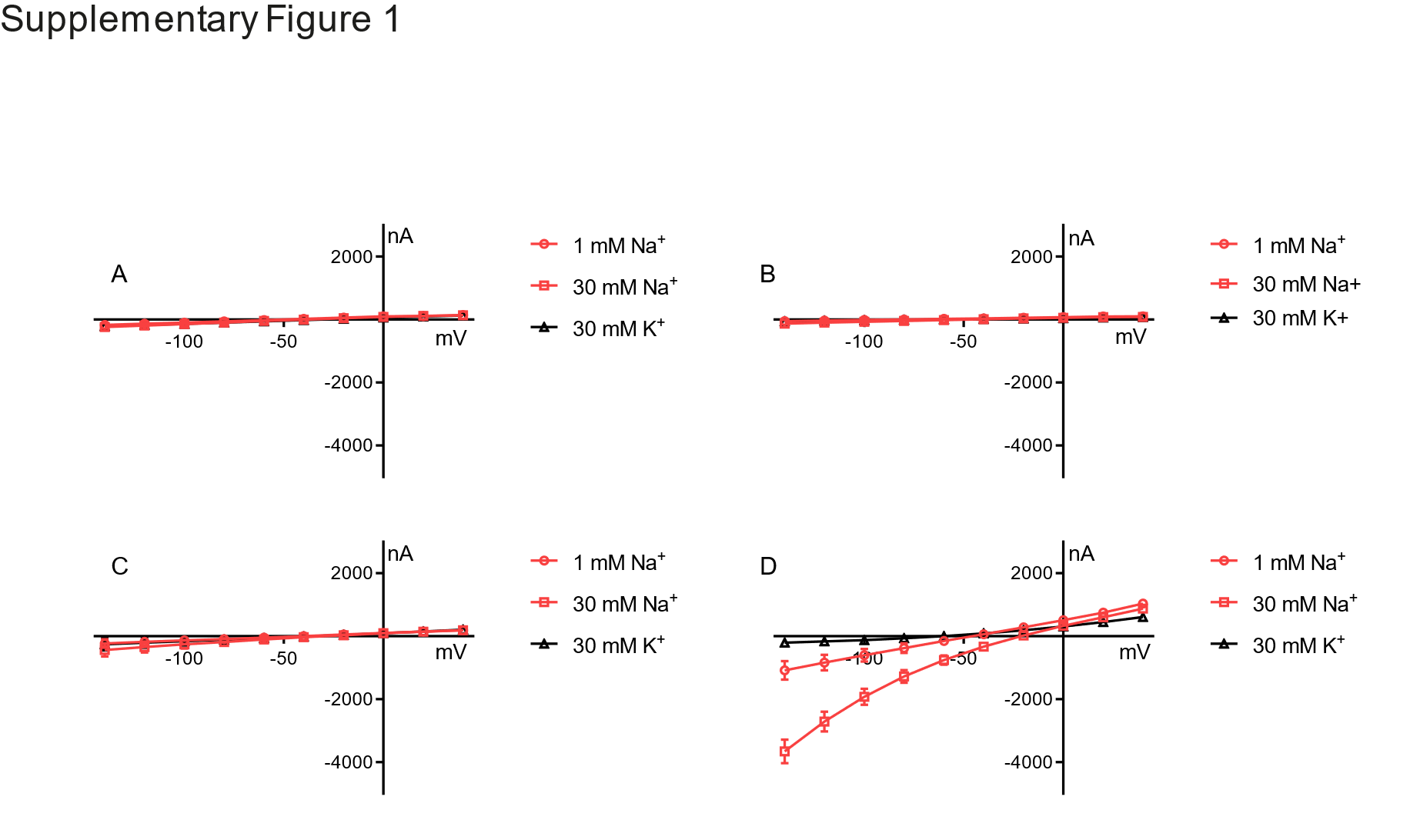

**Supplementary Figure 2:	Molecular models of HvHKT1;5_HAP3_L189_ and HvHKT1;5_HAP3_P189_ transporters in complex with Na^+^. a., b.** Cartoon representations of HvHKT1;5_HAP3_L189_ (left, grey) and HvHKT1;5_HAP3_P189_ (right, deep teal) with cylindrical α-helices illustrate 3D folds. Constrictions in selectivity filters are bound by four residues (cpk magenta sticks, regular types) that contain Na^+^ (violet spheres). Black arrows illustrate directional flows of Na^+^ that are likely to enter the permeation trajectory by-passing selectivity filter constrictions. Variant residues N57, V416, S438 and L189 (cpk sticks and dots) in HvHKT1;5_HAP3_l189_, and N57, V416 and S438 and P189 (cpk sticks and dots) in HvHKT1;5_HAP3_P189_ are indicated; the dots illustrate volumes of van der Waals radii. Four variations N57, V416 and S438 (regular types), and L189 or P189 (bold types) are shown in HvHKT1;5_HAP3_ proteins, from which P189 is deemed to be critical for protein structure that underlies function. **c., d.** Detailed views of α-helices, which neighbour constrictions of selectivity filters containing Na^+^ that are crucial for permeation function. Na^+^ (violet spheres) are located near the selectivity filter residues S76, G232, G351, G451 (cpk magenta sticks) in HvHKT1;5_HAP3_ structures. In each protein, polar contacts of L189 and P189 (shown in cpk sticks and dots), that are positioned on α-helix 4, are indicated by dashed lines at separations between 2.5 Å and 3.1 Å. Notably, leucine or proline residues in HvHKT1;5_HAP3_ structures effect packing angles of bordering α-helices 4 and 5. This packing angle between α-helices 4 and 5 in HvHKT1;5_HAP3_P189_ is more obtuse (two black arrows pointing to each other).

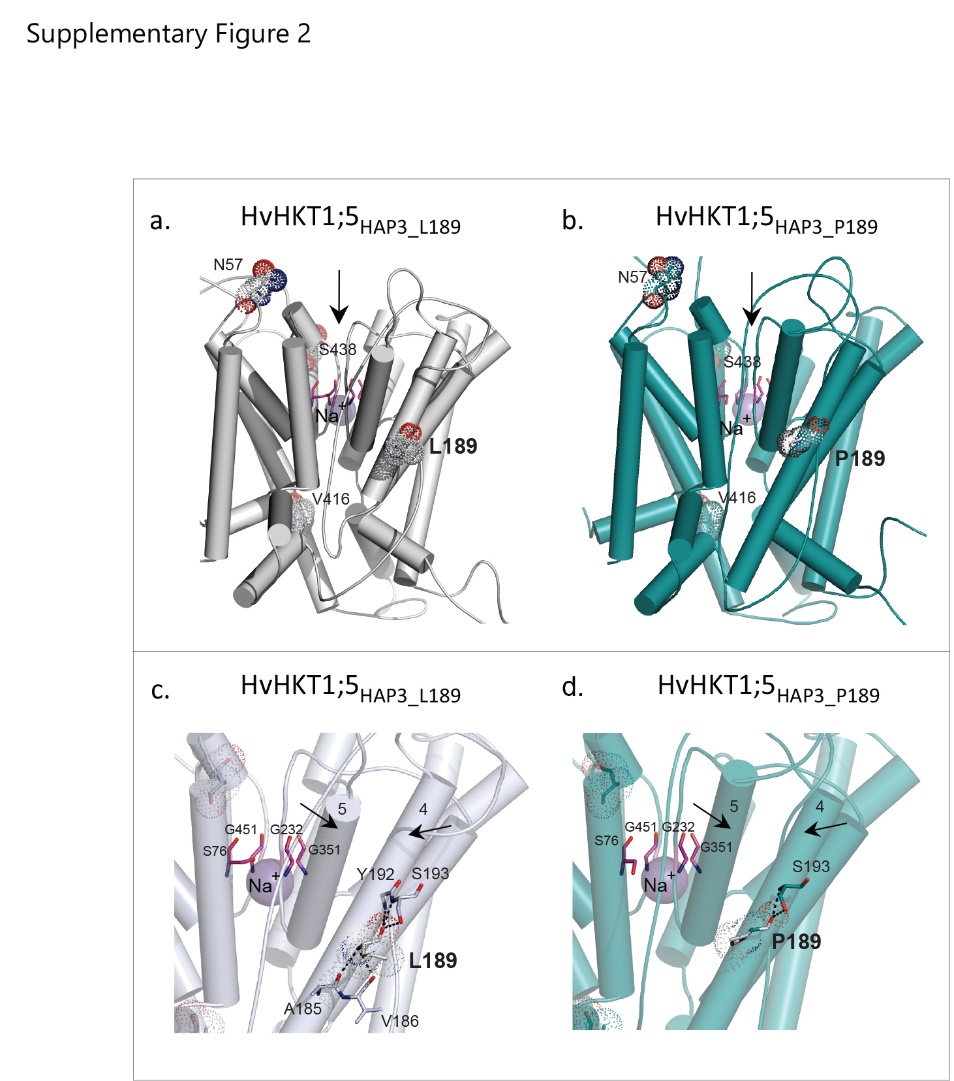

Supplementary Figure 2 Methods.

The most suitable template for HvHKT1;5 transporter proteins was the *B. subtilis* KtrB K+ transporter (Protein Data Bank accession 4J7C, chain I) (Vieira-Pires *et al*., 2013) as previously identified (Xu *et al*., 2018). In KtrB, K^+^ was substituted by Na^+^ during modelling of all HKT1;5 proteins. 3D models of HvHKT1;5_HAP3_L189_ and HvHKT1;5_HAP3_P189_ in complex with Na+ were generated in Modeller 9v19 (Sali and Blundell, 1993) as described previously (Cotsaftis *et al*., 2012, Waters *et al*., 2013) incorporating Na^+^ ionic radii (Xu *et al*., 2018) taken from the CHARMM force field (Brooks *et al*., 2009), on the Linux station running the Ubuntu 12.04 operating system. Best scoring models (from an ensemble of 50) were selected based on the combination of Modeller Objective Function (Shen and Sali, 2006), Discrete Optimised Protein Energy term (Eswar *et al*., 2008), PROCHECK (Laskowski *et al*., 1993), ProSa 2003 (Sippl, 1993) and FoldX (Schymkowitz *et al*., 2005). Structural images were generated in the PyMOL Molecular Graphics System V1.8.2.0 (Schrődinger LLC, Portland, OR, USA). Calculations of angles between selected α-helices in HvHKT1;5 models were executed in Chimera (Pettersen *et al*., 2004) and evaluations of differences (ΔΔG = ΔGmut-ΔGwt) in Gibbs free energies was performed with FoldX (Schymkowitz *et al*., 2005). Sequence conservation patterns were analysed with ConSurf (Landau *et al*., 2005; Celniker *et al*., 2013) based on 3D models of HvHKT1;5 transporters.

Evaluations of stereo-chemical parameters indicated that the template and HvHKT1;5 models had satisfactory parameters as indicated by Ramachandran plots with two residues positioned in disallowed regions, corresponding to 0.5% of all residues, except of G and P. Average G-factors (measures of correctness of dihedral angles and main-chain covalent bonds) of the template, and the HvHKT1;5_HAP3_L189_ and HvHKT1;5_HAP3_P189_ models, calculated by PROCHECK (0.06, -0.07 and -0.21, respectively), and ProSa 2003 z-scores (measures of Cβ-Cβ pair interactions of -9.0, -5.6 and -6.5, respectively), indicated that template and modelled structures had favourable conformational energies.

Results and Discussion

1.	Positional sequence identities between template and target sequences are in the twilight zone (20.6% and 20.4% between the template and HvHKT1;5_HAP3_L189_ and HvHKT1;5_HAP3_P189_ sequences, respectively), emphasising the difficulty of comparative modelling. This indicated that the attention must be paid to sequence alignments to be able to compare 3D models at the structural levels. Three types of alignments were generated, using Muscle (Edgar *et al*., 2004), MUSTER (Wu and Zhang, 2007) and LOMETS (Wu and Zhang, 2008) algorithms. Input alignments for 3D modelling were generated by the combination of all alignments and secondary structure elements analyses using PsiPred (Buchan *et al*., 2013), followed by manual adjustments to optimise positions of gaps in alignments.

2.	3D modelling revealed that overall 3D folds were similar, where selectivity filter constrictions carry one serine and three glycine residues, in accordance with their Na^+^ ion conductivity (Supplementary Figure 2).

3.	Detailed analysed of environments around α-helix 4 and α-helix 5 (two black arrows pointing to each other in Supplementary Figure 2) revealed that L189 in α-helix 4 of HvHKT1;5_HAP3_L189_ established four polar contacts at separations between 2.7 Å to 3.1 Å with A185, V186, Y192 and S193 neighbouring residues.

4.	These extensive polar contacts were not formed in the HvHKT1;5_HAP3_P189_ variant which only established two polar contacts at separations between 2.5 Å to 2.7 Å with S193. The lack of these cooperative binding networks in α-helix 4 around P189, and tight separations may impose severe structural rigidity on 3D folds HvHKT1;5_HAP3_P189_. These α-helices might no longer properly function in the structural and functional context to ensure Na^+^ ion conductance.

5.	In HvHKT1;5_HAP3_L189_, the packing angle between α-helix 4 and α-helix 5 (two black arrows pointing to each other in Supplementary Figure 2) is 9^o^ sharper compared to that in HvHKT1;5_HAP3_P189_ indicating that proline positions affect packing of α-helices in the specific 3D environments of HvHKT1;5 transporters. These changes in structural packing most likely contribute significantly to the structural rigidity and the lack of dynamics in 3D folds during transport.

6.	Evaluations of differences of Gibbs free energies of HvHKT1;5_HAP3_ transporters revealed that the L189P mutation was energetically highly unfavourable (highly destabilising), and that the reverse mutation (P189 into L189) restored 100% of this energy loss, as expected.

7.	In HvHKT1;5_HAP3_ transporters the differences in Gibbs free energies (ΔΔG) between the P189L variant and the reverse mutation (L189P) were mildly destabilising and similar in both directions, suggesting that the environment of P189 has somewhat adapted to its 3D fold, thus a low level of conductance of Na^+^ could be observed in HvHKT1;5_HAP3_P189__.

8.	In HvHKT1;5_HAP3_ we identified a positive correlation between structural characteristics of α-helix 4/α-helix 5 (trends in angles based on α-helical planes), differences in Gibbs free energies of forward (P189L) and reverse (L189P) mutations, and the ability to conduct Na^+^. This correlation shows that in barley HvHKT1;5 transporters, conservation and variability of specific residues reflect profoundly on the transport function.

9.	Sequence conservation patterns, based on 3D models of HvHKT1;5_HAP3_ using 368-370 sequences at sequence identities of 30% and higher (specifications: HMMMER homolog search algorithm, UNIREF-90 Protein database with the E-value cutoff of 1·10^-4^, Bayesian Model of substitution for proteins), revealed that the P189 variation occurred only in barley.

10.	There are four variations (N57, P189, V416 and S438) in HvHKT1;5_HAP3_ compared to HvHKT1;5_HAP1_ that represent 15 testable combinations. Not all of them could be tested for transport function. We suggest that three variations (N57, V416, S438) between HvHKT1;5_HAP3_ and HvHKT1;5_HAP1_ would have a lesser impact on Na^+^ conductivity. This is supported by conservation patterns analyses showing that these residues could be substituted by a variety of (mostly hydrophilic) residues, namely to R, S, A, P, G, L, H, D, Y, V, N, T, E, I (for N57), F, L, I, A, T, V (for V416), and N, T, Y, K, Q, S, R, H, A, P (for S438).

**Supplementary Figure 3: Influence of L189P polymorphism in HvHKT1;5 on grain K^+^ accumulation.** Mature grain K^+^ content after barley accessions were exposed to different levels of NaCl at the fourth leaf stage of development. White bars indicate 0mM of NaCl, light grey bars indicate 150mM NaCl, and dark grey indicates 250Mm NaCl added to plants.

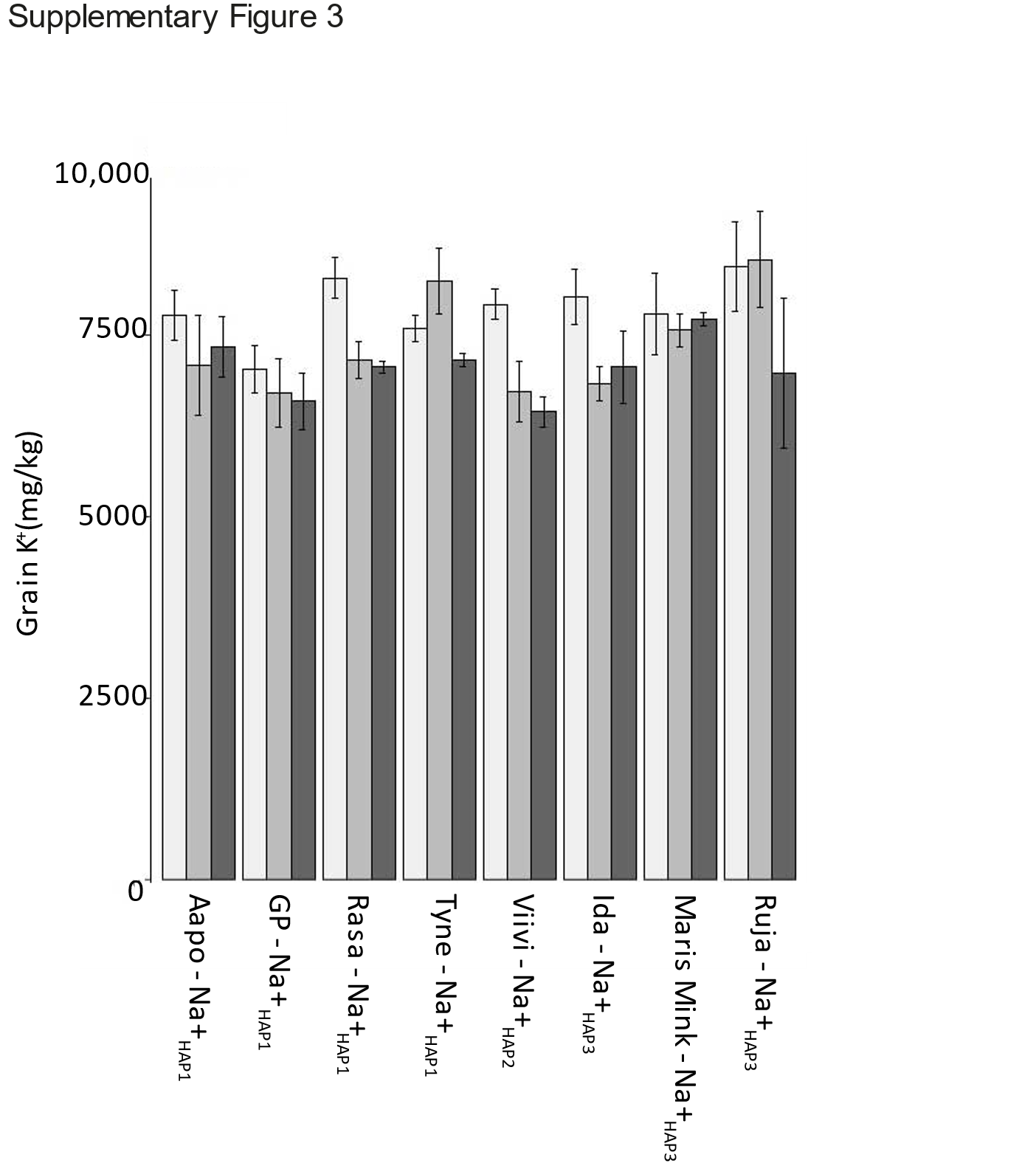

**Supplementary Figure 4: Influence of L189P polymorphism in HvHKT1;5 on shoot Na^+^**

**accumulation.** Fifth leaf Na^+^ content after barley accessions were exposed to different levels of

NaCl at the fourth leaf stage of development. White bars indicate 0mM of NaCl, light grey bars

indicate 150mM NaCl, and dark grey indicates 250Mm NaCl added to plants.

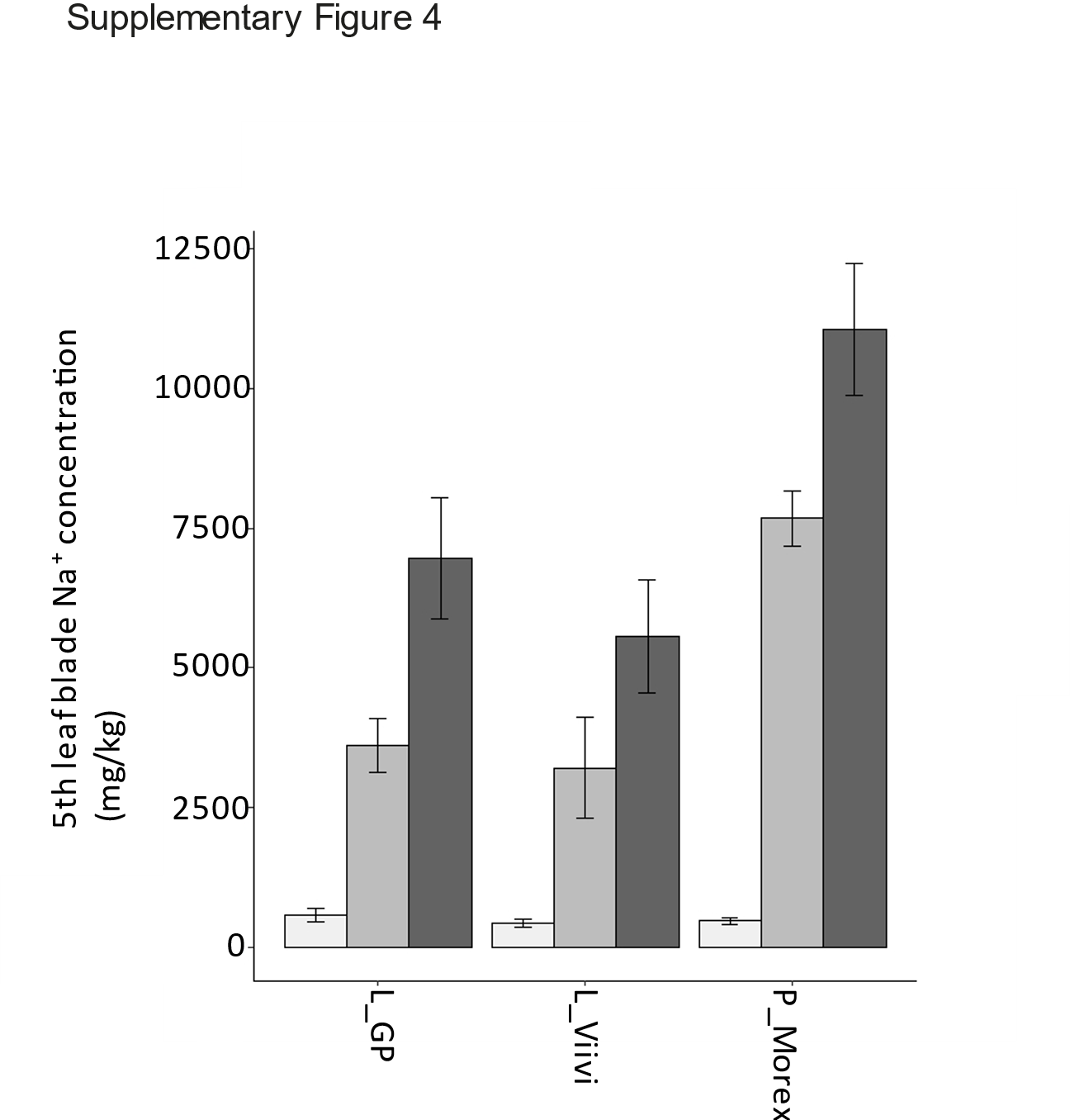

**Supplementary Figure 5:** Multiple alignment of species HKT orthologues

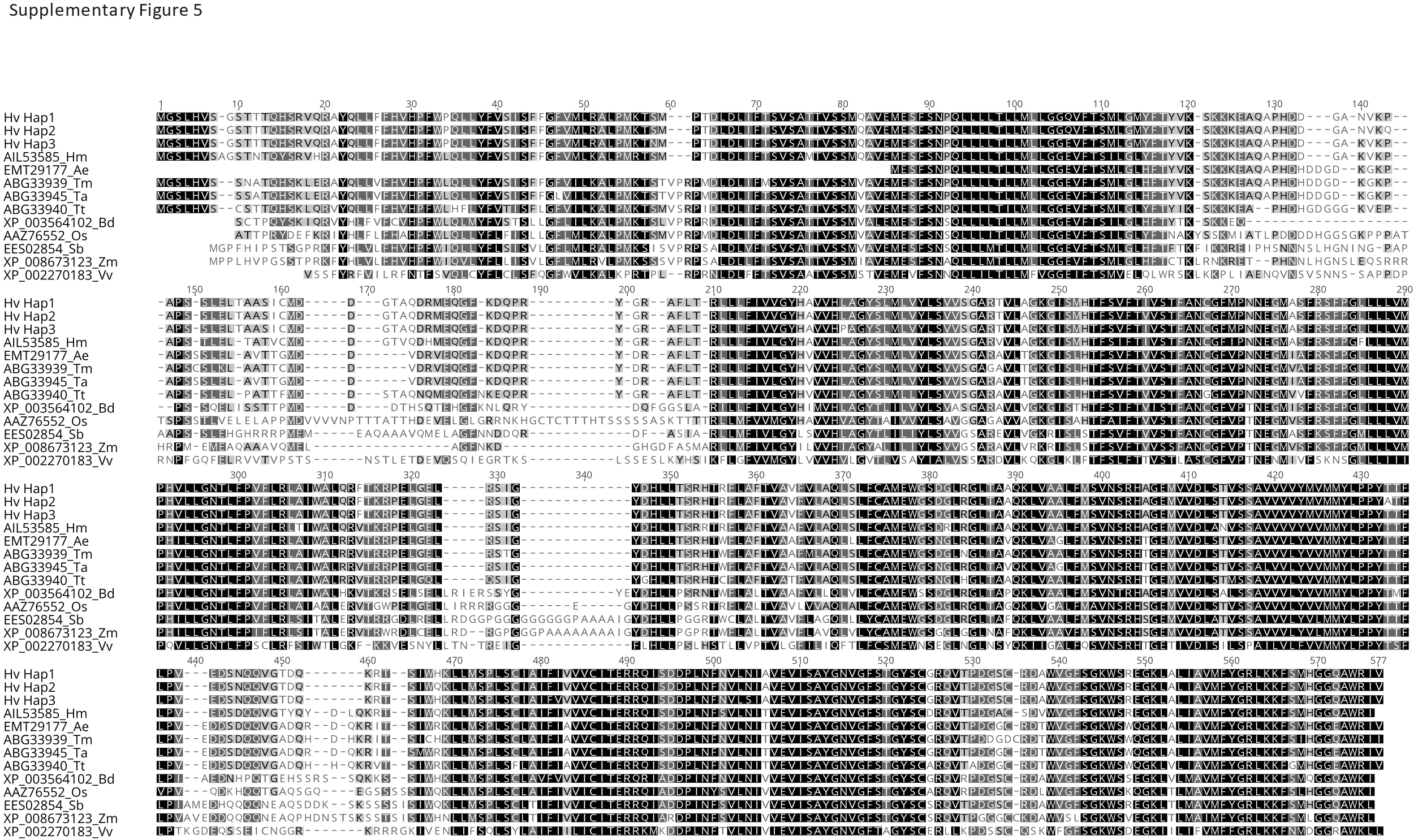

**Supplementary Figure 6: Geographical distribution of L189P in barley germplasm.**

**a.** Geographical distribution of L189P in *HvHKT1;5* in *H. spontaneum*. Location of accessions containing L189 in blue and 189P in red. **b**. Geographical distribution of L189P in *HvHKT1;5* in *H. vulgare landraces*. Location of accessions containing L189 in blue (Na^+^_HAP1,_ Na^+^_HAP2_) and 189P in red (Na^+^_HAP3_).

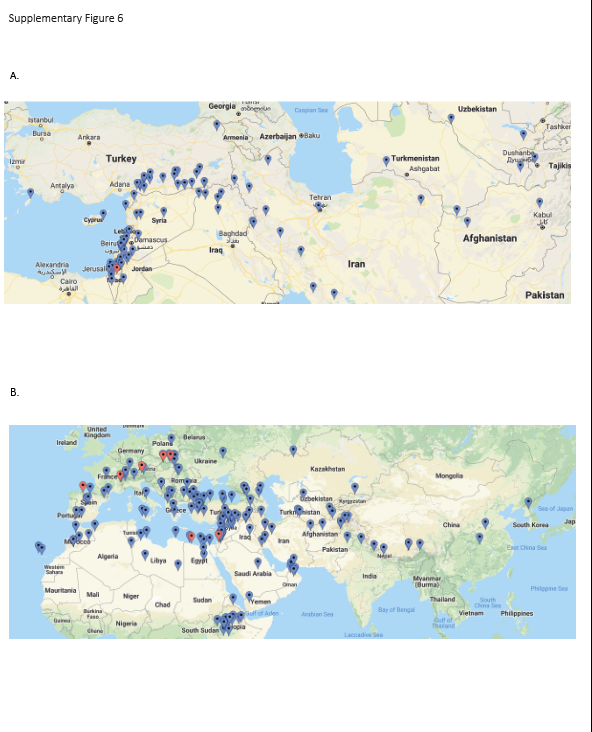

**Supplementary Figure 7.** Distribution of *HvHKT1;5* haplotypes in different genepools

**A.** Geographical distribution of L189P in *HvHKT1;5* in *H. vulgare landraces*. Location of accessions containing L189 in blue and 189P in red. **B**. Dendrogram using 4000 snps selected randomly from across the genome of H. *vulgare landraces,* accessions containing L189 are in blue and 189P in red.

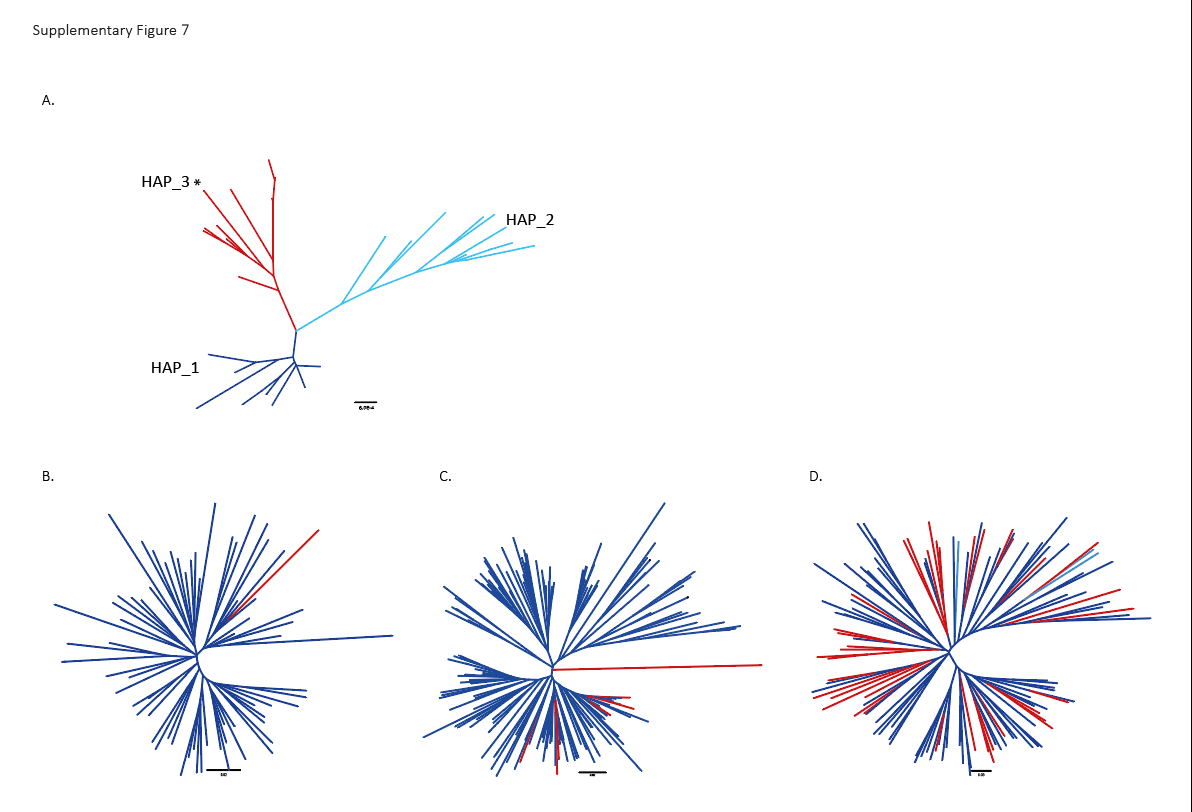

**Supplementary Tables**

***See associated excel sheets***

#### Supplementary dataset: Influence of growth in 0mM, 150mM and 250mM on a range of phenotypic traits

#### Biomass and Grain Data set sample numbers

Treatment

| Treatment | Biomass  Data  Freq | Grain  Data  Freq |
| --- | --- | --- |
| Control | 39 | 31 |
| 150 mM NaCl | 40 | 28 |
| 250 mM NaCl | 38 | 23 |

Allele, haplotype and line

| Allele | Haplotype | Line | Freq  Biomass Data | Freq  Grain Data |
| --- | --- | --- | --- | --- |
| L | Hap1 | Aapo | 15 | 11 |
| L | Hap1 | GP | 15 | 12 |
| L | Hap1 | Rasa | 15 | 8 |
| L | Hap1 | Tyne | 15 | 9 |
| L | Hap2 | Viivi | 15 | 10 |
| P | Hap3 | Ida | 14 | 10 |
| P | Hap3 | Maris_Mink | 15 | 13 |
| P | Hap3 | Ruja | 13 | 9 |

#### Shoot data set sample numbers

Treatment

| Treatment | Freq |
| --- | --- |
| Control | 21 |
| 150 mM NaCl | 20 |
| 250 mM NaCl | 20 |

Line by Allele

| Allele | Line | Freq |
| --- | --- | --- |
| L | Golden_promise | 20 |
| L | Viivi | 21 |
| P | Morex | 20 |

### Analysis of phenotypic traits

There is confounding between Allele (2 levels), Haplotype (3 levels) and Line (8 levels) as one particular line can only have a single allele and haplotype-so essentially there are only 8 combinations of allele, haplotype and line. When fitting a model it is therefore not possible to fit a crossed interaction between these 3 factors as many of the combinations do not exist within the data set. The treatment structure for these 3 factors needs to be nested: Allele/Haplotypes/Lines. So lines are within haplotypes which are within alleles.

The linear model approach presented aims to isolate whether the differences in trait are attributable to the alleles, haplotypes within alleles or lines within haplotypes within alleles or their interactions.

Salt treatment is also of interest, therefore a nested treatment structure between factors Allele, Haplotypes and Lines crossed with the salt Treatment can be used. The terms in the model are as follows:

Terms in the model

| Term | Interpretation | df |
| --- | --- | --- |
| (Intercept) | Overall mean | 1 |
| Allele | Allele | 1 |
| Treatment | Treatment | 2 |
| Allele:Haplotype | Haplotype | 1 |
| Allele:Treatment | Allele:Treatment | 2 |
| Allele:Haplotype:Line | Line | 5 |
| Allele:Haplotype:Treatment | Haplotype:Treatment | 2 |
| Allele:Haplotype:Line:Treatment | Line:Treatment | 10 |

For the shoot Na, only three lines were included in the experiment, therefore Haplotype and Line are completely confounded and only one can be included in the model. The terms in the model are as follows:

Terms in the model

| Term | Interpretation | df |
| --- | --- | --- |
| (Intercept) | Overall Mean | 1 |
| Allele | Allele | 1 |
| Treatment | Treatment | 2 |
| Allele:Haplotype | Haplotype | 1 |
| Allele:Treatment | Allele:Treatment | 2 |
| Allele:Haplotype:Treatment | Haplotype:Treatment | 2 |

Significance was tested at the 5% level. In general, terms in a model should be tested in a hierarchical manner, however, because of the confounding in this model there are exceptions to this. For example, the three-way interaction Allele:Haplotype:Treatment and the two way interaction Allele:Treatment can be examined simultaneously with the 4-way interaction. In addition, because of confounding there is an alternative interpretation of some terms. For example the three-way interaction between Allele, Haplotype and Line can be interpreted as a line effect and the two way interaction between Allele and Haplotype corresponds to haplotype effects. Because of the difficult in interpretation of relevant significant terms, in the Anova tables below, terms highlighted are the appropriate significant terms for interpretation.

ASReml-R was which uses a REML approach was used for analysis because of the unbalanced nature of the data. Anova tables presented are from ASReml. For some lower order terms predicted values were not available from ASReml due to missing treatment combinations. For these terms predicted values were obtained using a linear regression model in Genstat. For consistency, if lower order terms were significant, all predicted values available are from a linear regression model conducted in Genstat. Differences between treatment levels was determined by tukeys if the standard error of difference variance covariance matrix was available or By a bonferonni corrected LSD (least significant difference) if just the average standard error of difference was available.

#### Biomass

There is a significant three-way interaction between Allele, Haplotype and Line and therefore the trait differs between lines. There is a significant interaction between Allele and Haplotype (borderline) and there is also a main effect of Treatment and of Allele. The treatment effect is independent of allele, haplotype and line. All predicted values are from Genstat regression.

Anova Table: Biomass

| Term | Df | Sum of Sq | Wald statistic | Pr(Chisq) |
| --- | --- | --- | --- | --- |
| Allele | 1 | 17.3170719 | 10.3121835 | 0.0013215 |
| Treatment | 2 | 240.7291349 | 143.3523542 | 0.0000000 |
| Allele:Haplotype | 1 | 6.5975704 | 3.9288026 | 0.0474657 |
| Allele:Treatment | 2 | 3.3946952 | 2.0215150 | 0.3639432 |
| Allele:Haplotype:Line | 5 | 22.2350352 | 13.2407930 | 0.0212238 |
| Allele:Haplotype:Treatment | 2 | 0.2980298 | 0.1774745 | 0.9150860 |
| Allele:Haplotype:Line:Treatment | 10 | 17.4613580 | 10.3981048 | 0.4062873 |

Predicted Values (Allele by Haplotype by) Line: Biomass

| Allele | Haplotype | Line | Predicted value | Std error | Group* |
| --- | --- | --- | --- | --- | --- |
| L | Hap1 | Aapo | 7.518 | 0.33467 | abc |
| L | Hap1 | GP | 7.243 | 0.33467 | ab |
| L | Hap1 | Rasa | 6.823 | 0.33467 | a |
| L | Hap1 | Tyne | 6.688 | 0.33467 | a |
| L | Hap2 | Viivi | 7.807 | 0.33467 | abc |
| P | Hap3 | Ida | 7.125 | 0.34765 | ab |
| P | Hap3 | Maris_Mink | 8.195 | 0.33467 | bc |
| P | Hap3 | Ruja | 8.508 | 0.36082 | b |

*predicted mean values compared with bonferonni corrected LSD

Predicted Values Treatment: Biomass

| Treatment | Predicted value | Std error | Group* |
| --- | --- | --- | --- |
| Control | 7.518 | 0.2297 | a |
| 150 mM NaCl | 8.814 | 0.2290 | b |
| 250 mM NaCl | 5.423 | 0.2306 | c |

*predicted mean values compared with bonferonni corrected LSD

Predicted Values Haplotype: Biomass

| Allele | Haplotype | Predicted value | Std error | Group* |
| --- | --- | --- | --- | --- |
| L | Hap1 | 7.068 | 0.1673 | a |
| L | Hap2 | 7.807 | 0.3347 | b |
| P | Hap3 | 7.935 | 0.2005 | b |

*predicted mean values compared by bonferonni corrected LSD

Predicted Values Allele: Biomass

| Allele | Predicted value | Std error |
| --- | --- | --- |
| L | 7.111 | 0.2557 |
| P | 7.935 | 0.2557 |

#### Ear weight

There is a significant four-way interaction between Allele, Haplotype, Line and Treatment. Therefore the ear weight depends on the line and on the treatment.

Anova Table: Ear weight

| Term | Df | Sum of Sq | Wald statistic | Pr(Chisq) |
| --- | --- | --- | --- | --- |
| Allele | 1 | 0.5417924 | 0.2075091 | 0.6487268 |
| Treatment | 2 | 95.5848289 | 36.6094558 | 0.0000000 |
| Allele:Haplotype | 1 | 9.0859323 | 3.4799564 | 0.0621164 |
| Allele:Treatment | 2 | 10.9303008 | 4.1863585 | 0.1232945 |
| Allele:Haplotype:Line | 5 | 44.8262946 | 17.1686895 | 0.0041907 |
| Allele:Haplotype:Treatment | 2 | 5.4976098 | 2.1056114 | 0.3489573 |
| Allele:Haplotype:Line:Treatment | 10 | 65.9813122 | 25.2711645 | 0.0048545 |

Predicted Values (Allele by Haplotype by) Line by Treatment: Ear weight

| Allele | Haplotype | Line | Treatment | Predicted value | Std error | Group* |
| --- | --- | --- | --- | --- | --- | --- |
| L | Hap1 | Aapo | Control | 4.03220 | 0.7226248 | abcd |
| L | Hap1 | Aapo | 150 mM NaCl | 3.95380 | 0.7226248 | abcd |
| L | Hap1 | Aapo | 250 mM NaCl | 2.16000 | 0.7226248 | abc |
| L | Hap1 | GP | Control | 1.61380 | 0.7226248 | ab |
| L | Hap1 | GP | 150 mM NaCl | 5.43260 | 0.7226248 | bcd |
| L | Hap1 | GP | 250 mM NaCl | 2.52100 | 0.7226248 | abc |
| L | Hap1 | Rasa | Control | 3.64140 | 0.7226248 | abcd |
| L | Hap1 | Rasa | 150 mM NaCl | 5.45900 | 0.7226248 | cd |
| L | Hap1 | Rasa | 250 mM NaCl | 3.47240 | 0.7226248 | abcd |
| L | Hap1 | Tyne | Control | 6.37020 | 0.7226248 | d |
| L | Hap1 | Tyne | 150 mM NaCl | 6.42920 | 0.7226248 | d |
| L | Hap1 | Tyne | 250 mM NaCl | 2.75220 | 0.7226248 | abcd |
| L | Hap2 | Viivi | Control | 3.99380 | 0.7226248 | abcd |
| L | Hap2 | Viivi | 150 mM NaCl | 3.86900 | 0.7226248 | abcd |
| L | Hap2 | Viivi | 250 mM NaCl | 1.48620 | 0.7226248 | a |
| P | Hap3 | Ida | Control | 5.26840 | 0.7226248 | abcd |
| P | Hap3 | Ida | 150 mM NaCl | 3.04540 | 0.7226248 | abcd |
| P | Hap3 | Ida | 250 mM NaCl | 2.42975 | 0.8079191 | abcd |
| P | Hap3 | Maris_Mink | Control | 4.39020 | 0.7226248 | abcd |
| P | Hap3 | Maris_Mink | 150 mM NaCl | 5.74300 | 0.7226248 | cd |
| P | Hap3 | Maris_Mink | 250 mM NaCl | 2.30740 | 0.7226248 | abc |
| P | Hap3 | Ruja | Control | 3.22350 | 0.8079191 | abcd |
| P | Hap3 | Ruja | 150 mM NaCl | 3.17660 | 0.7226248 | abcd |
| P | Hap3 | Ruja | 250 mM NaCl | 2.97425 | 0.8079191 | abcd |

*predicted mean values compared with tukeys

#### Biomass Combined with ear weight

There is a significant four-way interaction between Allele, Haplotype, Line and Treatment. Therefore the biomass combined with ear weight depends on both the line and the treatment with a non-additive interaction.

Anova Table: Biomass Combined with ear weight

| Term | Df | Sum of Sq | Wald statistic | Pr(Chisq) |
| --- | --- | --- | --- | --- |
| Allele | 1 | 1.173277e+01 | 2.6159346 | 0.1057952 |
| Treatment | 2 | 6.374063e+02 | 142.1159035 | 0.0000000 |
| Allele:Haplotype | 1 | 1.986613e-01 | 0.0442935 | 0.8333086 |
| Allele:Treatment | 2 | 9.229523e+00 | 2.0578115 | 0.3573978 |
| Allele:Haplotype:Line | 5 | 3.632991e+01 | 8.1001046 | 0.1508043 |
| Allele:Haplotype:Treatment | 2 | 7.982469e+00 | 1.7797688 | 0.4107032 |
| Allele:Haplotype:Line:Treatment | 10 | 9.648986e+01 | 21.5133476 | 0.0177851 |

Predicted Values (Allele by Haplotype by) line by treatment: Biomass Combined with ear weight

| Allele | Haplotype | Line | Treatment | Predicted value | Std error | Group* |
| --- | --- | --- | --- | --- | --- | --- |
| L | Hap1 | Aapo | Control | 11.51860 | 0.9471131 | abcdef |
| L | Hap1 | Aapo | 150 mM NaCl | 12.97100 | 0.9471131 | bcdef |
| L | Hap1 | Aapo | 250 mM NaCl | 8.13340 | 0.9471131 | ab |
| L | Hap1 | GP | Control | 8.80900 | 0.9471131 | abcd |
| L | Hap1 | GP | 150 mM NaCl | 14.23520 | 0.9471131 | ef |
| L | Hap1 | GP | 250 mM NaCl | 8.17160 | 0.9471131 | ab |
| L | Hap1 | Rasa | Control | 10.81800 | 0.9471131 | abcde |
| L | Hap1 | Rasa | 150 mM NaCl | 13.76160 | 0.9471131 | def |
| L | Hap1 | Rasa | 250 mM NaCl | 8.37360 | 0.9471131 | ab |
| L | Hap1 | Tyne | Control | 13.49560 | 0.9471131 | cdef |
| L | Hap1 | Tyne | 150 mM NaCl | 14.54280 | 0.9471131 | ef |
| L | Hap1 | Tyne | 250 mM NaCl | 7.49040 | 0.9471131 | a |
| L | Hap2 | Viivi | Control | 12.15540 | 0.9471131 | abcdef |
| L | Hap2 | Viivi | 150 mM NaCl | 12.96200 | 0.9471131 | bcdef |
| L | Hap2 | Viivi | 250 mM NaCl | 7.57680 | 0.9471131 | a |
| P | Hap3 | Ida | Control | 13.04420 | 0.9471131 | bcdef |
| P | Hap3 | Ida | 150 mM NaCl | 11.01880 | 0.9471131 | abcde |
| P | Hap3 | Ida | 250 mM NaCl | 7.99300 | 1.0589046 | ab |
| P | Hap3 | Maris_Mink | Control | 12.35120 | 0.9471131 | abcdef |
| P | Hap3 | Maris_Mink | 150 mM NaCl | 16.14020 | 0.9471131 | f |
| P | Hap3 | Maris_Mink | 250 mM NaCl | 8.42520 | 0.9471131 | ab |
| P | Hap3 | Ruja | Control | 12.69000 | 1.0589046 | abcdef |
| P | Hap3 | Ruja | 150 mM NaCl | 13.86180 | 0.9471131 | ef |
| P | Hap3 | Ruja | 250 mM NaCl | 8.20725 | 1.0589046 | abc |

*predicted mean values compared with tukeys

#### TGW

There is a significant three-way interaction between Allele, Haplotype and Line and between Allele and Treatment.

Anova Table: TGW

| Term | Df | Sum of Sq | Wald statistic | Pr(Chisq) |
| --- | --- | --- | --- | --- |
| Allele | 1 | 10.34065 | 0.1274180 | 0.7211243 |
| Treatment | 2 | 558.85998 | 6.8863022 | 0.0319638 |
| Allele:Haplotype | 1 | 126.85402 | 1.5631019 | 0.2112116 |
| Allele:Treatment | 2 | 986.99781 | 12.1618390 | 0.0022861 |
| Allele:Haplotype:Line | 5 | 1190.09690 | 14.6644368 | 0.0118966 |
| Allele:Haplotype:Treatment | 2 | 21.45482 | 0.2643674 | 0.8761800 |
| Allele:Haplotype:Line:Treatment | 10 | 1202.71240 | 14.8198857 | 0.1387678 |

Predicted Values (Allele by Haplotype by) Line: TGW

| Allele | Haplotype | Line | Predicted value | Std error | Group* |
| --- | --- | --- | --- | --- | --- |
| L | Hap1 | Aapo | 40.433 | 2.3265 | b |
| L | Hap1 | GP | 30.658 | 2.3265 | a |
| L | Hap1 | Rasa | 38.521 | 2.3265 | b |
| L | Hap1 | Tyne | 34.419 | 2.3265 | ab |
| L | Hap2 | Viivi | 39.285 | 2.3265 | b |
| P | Hap3 | Ida | 36.752 | 2.4168 | ab |
| P | Hap3 | Maris_Mink | 32.606 | 2.3265 | ab |
| P | Hap3 | Ruja | 39.577 | 2.5083 | b |

*predicted mean values compared with corrected bonferonni LSD

Predicted Values Allele by Treatment: TGW

| Allele | Treatment | Predicted value | Std error | Group* |
| --- | --- | --- | --- | --- |
| L | Control | 33.13 | 1.911 | a |
| L | 150 mM NaCl | 42.02 | 1.911 | a |
| L | 250 mM NaCl | 33.23 | 1.911 | a |
| P | Control | 35.69 | 2.412 | a |
| P | 150 mM NaCl | 33.96 | 2.330 | a |
| P | 250 mM NaCl | 38.91 | 2.504 | a |

*predicted mean values compared with bonferonni corrected LSD

#### Area

There is a significant three-way interaction between Allele, Haplotype and Line and the two-way interaction between Allele and Treatment effect and between Allele and Haplotype.

Anova Table: Area Anova Table: Area

| Term | Df | Sum of Sq | Wald statistic | Pr(Chisq) |
| --- | --- | --- | --- | --- |
| Allele | 1 | 3.895407e-01 | 8.517500e-02 | 0.7704031 |
| Treatment | 2 | 9.862775e+00 | 2.156545e+00 | 0.3401826 |
| Allele:Haplotype | 1 | 1.276616e+02 | 2.791386e+01 | 0.0000001 |
| Allele:Treatment | 2 | 3.302763e+01 | 7.221657e+00 | 0.0270294 |
| Allele:Haplotype:Line | 5 | 9.551324e+01 | 2.088445e+01 | 0.0008518 |
| Allele:Haplotype:Treatment | 2 | 2.041867e+00 | 4.464644e-01 | 0.7999291 |
| Allele:Haplotype:Line:Treatment | 10 | 5.998659e+01 | 1.311637e+01 | 0.2172380 |

Predicted Values (Allele by Haplotype by) Line: Area

| Allele | Haplotype | Line | Predicted value | Std error | Group* |
| --- | --- | --- | --- | --- | --- |
| L | Hap1 | Aapo | 22.604 | 0.55229 | cd |
| L | Hap1 | GP | 20.159 | 0.55229 | a |
| L | Hap1 | Rasa | 21.296 | 0.55229 | abc |
| L | Hap1 | Tyne | 20.097 | 0.55229 | a |
| L | Hap2 | Viivi | 24.292 | 0.55229 | d |
| P | Hap3 | Ida | 22.180 | 0.57372 | bc |
| P | Hap3 | Maris_Mink | 20.680 | 0.55229 | ab |
| P | Hap3 | Ruja | 22.797 | 0.59545 | cd |

*predicted mean values compared with bonferonni corrected LSD

Predicted Values Allele by Treatment: Area

| Allele | Treatment | Predicted value | Std error | Group* |
| --- | --- | --- | --- | --- |
| L | Control | 20.84 | 0.4536 | a |
| L | 150 mM NaCl | 22.20 | 0.4536 | a |
| L | 250 mM NaCl | 20.61 | 0.4536 | a |
| P | Control | 21.67 | 0.5726 | a |
| P | 150 mM NaCl | 21.31 | 0.5531 | a |
| P | 250 mM NaCl | 22.56 | 0.5943 | a |

*predicted mean values compared with bonferonni corrected LSD

Predicted Values (Allele by) Haplotype: Area

| Allele | Haplotype | Predicted value | Std error | Group* |
| --- | --- | --- | --- | --- |
| L | Hap1 | 21.04 | 0.2761 | a |
| L | Hap2 | 24.29 | 0.5523 | b |
| P | Hap3 | 21.84 | 0.3308 | a |

*predicted mean values compared with bonferonni corrected LSD

#### Width

There is a significant interaction between allele and treatment.

Anova Table: Width

| Term | Df | Sum of Sq | Wald statistic | Pr(Chisq) |
| --- | --- | --- | --- | --- |
| Allele | 1 | 0.0265934 | 3.090325e-01 | 0.5782745 |
| Treatment | 2 | 0.1189233 | 1.381965e+00 | 0.5010836 |
| Allele:Haplotype | 1 | 0.0833333 | 9.683869e-01 | 0.3250828 |
| Allele:Treatment | 2 | 0.5880284 | 6.833268e+00 | 0.0328227 |
| Allele:Haplotype:Line | 5 | 0.8446874 | 9.815810e+00 | 0.0806256 |
| Allele:Haplotype:Treatment | 2 | 0.0330667 | 3.842559e-01 | 0.8252013 |
| Allele:Haplotype:Line:Treatment | 10 | 1.2746752 | 1.481255e+01 | 0.1390471 |

Predicted Values Allele by Treatment: Width

| Allele | Treatment | Predicted value | Std error | Group* |
| --- | --- | --- | --- | --- |
| L | Control | 3.136 | 0.06222 | a |
| L | 150 mM NaCl | 3.311 | 0.06222 | a |
| L | 250 mM NaCl | 3.098 | 0.06222 | a |
| P | Control | 3.194 | 0.07854 | a |
| P | 150 mM NaCl | 3.083 | 0.07587 | a |
| P | 250 mM NaCl | 3.230 | 0.08153 | a |

*predicted mean values compared with bonferonni corrected LSD

#### Length

There is significant three-way interactions between Allele, Haplotype and Treatment and between Allele, Haplotype and Line.

Anova Table: Length

| Term | Df | Sum of Sq | Wald statistic | Pr(Chisq) |
| --- | --- | --- | --- | --- |
| Allele | 1 | 0.1047680 | 6.900687e-01 | 0.4061410 |
| Treatment | 2 | 0.0501037 | 3.300145e-01 | 0.8478876 |
| Allele:Haplotype | 1 | 13.0208333 | 8.576348e+01 | 0.0000000 |
| Allele:Treatment | 2 | 0.1207161 | 7.951131e-01 | 0.6719599 |
| Allele:Haplotype:Line | 5 | 3.1422507 | 2.069686e+01 | 0.0009241 |
| Allele:Haplotype:Treatment | 2 | 0.9760667 | 6.428995e+00 | 0.0401755 |
| Allele:Haplotype:Line:Treatment | 10 | 0.8202914 | 5.402960e+00 | 0.8626875 |

Predicted Values (Allele by Haplotype by) Line: Length

| Allele | Haplotype | Line | Predicted value | Std error | Group* |
| --- | --- | --- | --- | --- | --- |
| L | Hap1 | Aapo | 8.768 | 0.10063 | c |
| L | Hap1 | GP | 8.339 | 0.10063 | a |
| L | Hap1 | Rasa | 8.389 | 0.10063 | ab |
| L | Hap1 | Tyne | 8.236 | 0.10063 | a |
| L | Hap2 | Viivi | 9.470 | 0.10063 | d |
| P | Hap3 | Ida | 8.864 | 0.10453 | c |
| P | Hap3 | Maris_Mink | 8.565 | 0.10063 | abc |
| P | Hap3 | Ruja | 8.683 | 0.10849 | abc |

*predicted mean values compared with bonferonni corrected LSD

Predicted Values (Allele by) Haplotype by Treatment: Length

| Allele | Haplotype | Treatment | Predicted value | Std error | Group |
| --- | --- | --- | --- | --- | --- |
| L | Hap1 | Control | 8.460 | 0.08713 | a |
| L | Hap1 | 150 mM NaCl | 8.500 | 0.08713 | a |
| L | Hap1 | 250 mM NaCl | 8.335 | 0.08713 | a |
| L | Hap2 | Control | 9.300 | 0.17425 | b |
| L | Hap2 | 150 mM NaCl | 9.340 | 0.17425 | b |
| L | Hap2 | 250 mM NaCl | 9.780 | 0.17425 | b |
| P | Hap3 | Control | 8.634 | 0.10432 | a |
| P | Hap3 | 150 mM NaCl | 8.687 | 0.10078 | a |
| P | Hap3 | 250 mM NaCl | 8.786 | 0.10829 | a |

*predicted mean values compared with bonferonni corrected LSD

#### Grain Na

A square root transformations for Grain Na was necessary in order to meet model assumptions. There is a significant four way interaction between Allele, Haplotype, Line and treatment. Therefore the Grain Na changes depending on the line and treatment. There is also a significant two-way interaction between Allele and Treatment and between Allele and Haplotype

Anova Table: Grain Na

| Term | Df | Sum of Sq | Wald statistic | Pr(Chisq) |
| --- | --- | --- | --- | --- |
| Allele | 1 | 5801.32847 | 134.599678 | 0.0000000 |
| Treatment | 2 | 1293.23865 | 30.005111 | 0.0000003 |
| Allele:Haplotype | 1 | 213.27734 | 4.948360 | 0.0261154 |
| Allele:Treatment | 2 | 1034.18945 | 23.994774 | 0.0000062 |
| Allele:Haplotype:Line | 5 | 2587.21671 | 60.027378 | 0.0000000 |
| Allele:Haplotype:Treatment | 2 | 50.30663 | 1.167191 | 0.5578890 |
| Allele:Haplotype:Line:Treatment | 10 | 1549.98297 | 35.961971 | 0.0000855 |

Back Transformed Predicted Values for (Allele by Haplotype by) Line by treatment: Grain Na

| Allele | Haplotype | Line | Treatment | Predicted value | Std error | groups | Back transformed predicted value | Back transformed Std error |
| --- | --- | --- | --- | --- | --- | --- | --- | --- |
| L | Hap1 | Aapo | Control | 14.48421 | 2.936005 | ab | 209.7922 | 85.05139 |
| L | Hap1 | Aapo | 150 mM NaCl | 18.68596 | 3.790366 | ab | 349.1649 | 141.65320 |
| L | Hap1 | Aapo | 250 mM NaCl | 17.69716 | 3.790366 | ab | 313.1894 | 134.15740 |
| L | Hap1 | GP | Control | 17.31248 | 2.936005 | ab | 299.7220 | 101.65905 |
| L | Hap1 | GP | 150 mM NaCl | 20.73275 | 3.282553 | ab | 429.8469 | 136.11270 |
| L | Hap1 | GP | 250 mM NaCl | 23.69988 | 3.790366 | ab | 561.6842 | 179.66240 |
| L | Hap1 | Rasa | Control | 15.40053 | 3.790366 | ab | 237.1763 | 116.74727 |
| L | Hap1 | Rasa | 150 mM NaCl | 15.34165 | 3.790366 | ab | 235.3663 | 116.30093 |
| L | Hap1 | Rasa | 250 mM NaCl | 13.27139 | 4.642231 | ab | 176.1297 | 123.21767 |
| L | Hap1 | Tyne | Control | 15.97358 | 3.282553 | ab | 255.1552 | 104.86823 |
| L | Hap1 | Tyne | 150 mM NaCl | 18.89766 | 3.790366 | ab | 357.1216 | 143.25809 |
| L | Hap1 | Tyne | 250 mM NaCl | 24.83871 | 4.642231 | ab | 616.9616 | 230.61405 |
| L | Hap2 | Viivi | Control | 13.93784 | 3.790366 | ab | 194.2633 | 105.65900 |
| L | Hap2 | Viivi | 150 mM NaCl | 10.49707 | 3.790366 | a | 110.1884 | 79.57545 |
| L | Hap2 | Viivi | 250 mM NaCl | 16.06363 | 3.282553 | ab | 258.0403 | 105.45944 |
| P | Hap3 | Ida | Control | 22.64528 | 3.790366 | ab | 512.8089 | 171.66780 |
| P | Hap3 | Ida | 150 mM NaCl | 31.14192 | 3.282553 | b | 969.8190 | 204.44997 |
| P | Hap3 | Ida | 250 mM NaCl | 27.02554 | 3.790366 | ab | 730.3798 | 204.87335 |
| P | Hap3 | Marris Mink | Control | 22.82041 | 2.936005 | ab | 520.7712 | 134.00166 |
| P | Hap3 | Marris Mink | 150 mM NaCl | 53.42992 | 2.936005 | c | 2854.7563 | 313.74097 |
| P | Hap3 | Marris Mink | 250 mM NaCl | 63.33185 | 3.790366 | c | 4010.9238 | 480.10175 |
| P | Hap3 | Ruja | Control | 21.99550 | 3.790366 | ab | 483.8020 | 166.74196 |
| P | Hap3 | Ruja | 150 mM NaCl | 30.50674 | 3.790366 | ab | 930.6611 | 231.26338 |
| P | Hap3 | Ruja | 250 mM NaCl | 30.88116 | 3.790366 | ab | 953.6458 | 234.10174 |

*predicted mean values on transformed scale compared with tukeys

Back Transformed Predicted Values for (Allele by) Treatment: Grain Na

| Allele | Treatment | Predicted value | Std error | Group* | Back transformed predicted value | Back transformed  Std error |
| --- | --- | --- | --- | --- | --- | --- |
| L | Control | 15.74 | 1.519 | a | 247.75 | 47.82 |
| L | 150 mM NaCl | 18.20 | 1.739 | a | 331.24 | 63.30 |
| L | 250 mM NaCl | 19.98 | 1.969 | a | 399.20 | 78.68 |
| P | Control | 22.53 | 1.991 | a | 507.60 | 89.71 |
| P | 150 mM NaCl | 40.02 | 1.900 | b | 1601.60 | 152.08 |
| P | 250 mM NaCl | 42.86 | 2.216 | b | 1836.98 | 189.96 |

* predicted mean values on transformed scale compared with bonferonni corrected LSD

Back Transformed Predicted Values for (Allele by) Haplotype: Grain Na

| Allele | Haplotype | Predicted value | Std error | Group* | Back transformed predicted value | Back transformed Std error |
| --- | --- | --- | --- | --- | --- | --- |
| L | Hap1 | 18.04 | 1.0472 | a | 325.44 | 37.78 |
| L | Hap2 | 13.36 | 2.1392 | b | 178.49 | 57.16 |
| P | Hap3 | 34.20 | 1.1720 | c | 1169.64 | 80.16 |

*predicted mean values on transformed scale compared with bonferonni corrected LSD

#### Grain K

There is a significant treatment effect. Allele, Haplotype and Line are not significant.

Anova Table: Grain K

| Term | Df | Sum of Sq | Wald statistic | Pr(Chisq) |
| --- | --- | --- | --- | --- |
| Allele | 1 | 2.639048e+06 | 2.4878311 | 0.1147297 |
| Treatment | 2 | 8.649643e+06 | 8.1540218 | 0.0169581 |
| Allele:Haplotype | 1 | 5.751935e+05 | 0.5422351 | 0.4615079 |
| Allele:Treatment | 2 | 5.369355e+03 | 0.0050617 | 0.9974724 |
| Allele:Haplotype:Line | 5 | 7.656622e+06 | 7.2179002 | 0.2049326 |
| Allele:Haplotype:Treatment | 2 | 1.365310e+06 | 1.2870778 | 0.5254297 |
| Allele:Haplotype:Line:Treatment | 10 | 7.635811e+06 | 7.1982817 | 0.7066027 |

Predicted Values treatment: Grain K

| Treatment | Predicted value | Std error | Group* |
| --- | --- | --- | --- |
| Control | 7751 | 192.7 | a |
| 150 mM NaCl | 7329 | 213.7 | ab |
| 250 mM NaCl | 7077 | 243.0 | b |

* predicted mean values of salt treatment compared to control with LSD.

#### Shoot Na

A square root transformations for Grain Na was necessary in order to meet model assumptions. There is a significant two-way interaction between Allele and Treatment. Therefore, the response differs depending on both the allele and treatment.

Anova Table: Shoot Na

| Term | Df | Sum of Sq | Wald statistic | Pr(Chisq) |
| --- | --- | --- | --- | --- |
| Allele | 1 | 7056.94885 | 63.6587791 | 0.0000000 |
| Treatment | 2 | 44749.94004 | 403.6768024 | 0.0000000 |
| Allele:Haplotype | 1 | 280.59661 | 2.5311842 | 0.1116164 |
| Allele:Treatment | 2 | 2462.70965 | 22.2154209 | 0.0000150 |
| Allele:Haplotype:Treatment | 2 | 65.64216 | 0.5921397 | 0.7437355 |

Back Transformed Predicted Values for Allele by Treatment: Shoot Na

| Allele | Treatment | Predicted value | Std error | Group* | Back transformed predicted value | Back transformed  Std error |
| --- | --- | --- | --- | --- | --- | --- |
| L | Control | 21.77 | 2.721 | a | 473.93 | 118.47 |
| L | 150 mM NaCl | 57.01 | 2.924 | b | 3250.14 | 333.39 |
| L | 250 mM NaCl | 77.64 | 2.935 | c | 6027.97 | 455.75 |
| P | Control | 21.51 | 4.298 | a | 462.68 | 184.90 |
| P | 150 mM NaCl | 87.44 | 3.980 | c | 7645.75 | 696.02 |
| P | 250 mM NaCl | 104.56 | 3.980 | d | 10932.79 | 832.30 |

*predicted mean values on transformed scale compared with bonferonni corrected LSD
